# Selective Aurora A-TPX2 interaction inhibitors have *in vivo* efficacy as targeted anti-mitotic agents

**DOI:** 10.1101/2023.03.22.533679

**Authors:** Simon R. Stockwell, Duncan E. Scott, Gerhard Fischer, Estrella Guarino, Timothy P. C. Rooney, Tzu-Shean Feng, Tommaso Moschetti, Rajavel Srinivasan, Esther Alza, Alice Asteian, Claudio Dagostin, Anna Alcaide, Mathieu Rocaboy, Beata Blaszczyk, Alicia Higueruelo, Xuelu Wang, Maxim Rossmann, Trevor R. Perrior, Tom L. Blundell, David R. Spring, Grahame McKenzie, Chris Abell, John Skidmore, Ashok R. Venkitaraman, Marko Hyvönen

## Abstract

The protein kinase Aurora A, and its close relative, Aurora B, regulate human cell division. Aurora A is frequently overexpressed in cancers of the breast, ovary, pancreas and blood, provoking genome instability and resistance to anti-mitotic chemotherapy. Intracellular localization and enzymatic activity of Aurora A are regulated by its interaction with the spindle assembly factor TPX2. Here, we have used fragment-based, structure-guided lead discovery to develop small-molecule inhibitors of the Aurora A-TPX2 protein-protein interaction (PPI). These compounds act by novel mechanism compared to existing Aurora A inhibitors and they are highly specific to Aurora A over Aurora B. Our biophysically, structurally and phenotypically validated lead compound, **CAM2602**, exhibits oral bioavailability, favourable pharmacokinetics, pharmacodynamic biomarker modulation, and arrest of growth in tumour xenografts. Consistent with our original finding that Aurora A overexpression drives taxane-resistance in cancer cells, inhibition of Aurora A-TPX2 PPI synergizes with paclitaxel to suppress the outgrowth of pancreatic cancer cells. Our results provide a blueprint for targeting the Aurora A-TPX2 PPI for cancer therapy and suggest a promising clinical utility for this mode of action.

## Introduction

Aurora A is a serine/threonine protein kinase that plays an important role in controlling early stages of mitosis, including centrosome maturation and separation, mitotic entry, and bipolar spindle formation^[1,2]^. Aurora A may be upregulated in cancer cells as a consequence of chromosome rearrangements, aberrant gene expression, or through protein stabilisation. Aurora A overexpression is a common feature of several cancers including ovarian, prostate, pancreas and breast, and has been linked to poor treatment outcome^[3–5]^. Disruption of the spindle assembly checkpoint due to Aurora A overexpression promotes tumourigenesis via chromosomal instability and aneuploidy^[3,5–7]^. Conversely, genomically-unstable cancer cells may become critically reliant on Aurora A function^[8,9]^. Androgen-receptor positive models of castration-resistant prostate cancer also show significant sensitivity to Aurora A inhibition^[10]^. Furthermore, non-genetic elevation of Aurora A levels is reported to drive resistance to current generation EGFR inhibitors in non-small cell lung cancer models^[11]^ and tumour resistance to taxanes is a further consequence of aberrant expression^[12,13]^. Aurora A inhibitors are also increasingly finding use against AML and related leukaemias^[14–16]^. Consequently, the cancer therapeutic promise of an effective inhibitor of Aurora A is of much interest and the focus of multiple drug discovery studies^[17–19]^.

Targeting protein for *Xenopus* kinesin-like protein 2 (TPX2) is a spindle assembly factor essential for mitotic spindle organisation, maintaining spindle-pole integrity and microtubule nucleation^[20]^. Its interaction with Aurora A mediates localisation of Aurora A to spindle microtubules^[21]^, regulates Aurora A kinase activity by stabilization of the active protein^[22,23]^ and protects the activating Thr288 residue in the catalytic domain of Aurora A from the action of PP1 phosphatase^[24,25]^. Aurora A and TPX2 are frequently co-overexpressed in tumours^[26]^, therefore the association of Aurora A and TPX2 comprises a novel oncogenic unit that presents a promising target for cancer therapy^[1,22]^.

Significant effort has been applied to developing ATP-competitive inhibitors of the Aurora kinases and several have progressed to clinical trials ^[17,27,28]^. Reported Aurora A inhibitors bind to the highly conserved ATP-binding site of the kinase and consequently exhibit variable selectivity for Aurora A over related kinases, most notably Aurora B and Aurora C^[17,29]^. High similarity between Aurora A and Aurora B, especially in their catalytic sites^[30]^, makes it challenging to develop highly selective small molecule inhibitors for Aurora A. Alisertib (MLN8237, **Fig. 1A**)^[31]^, an Aurora A inhibitor in clinical trials, is reported to have a selectivity for Aurora A over Aurora B of approximately 200-fold^[32]^, although work using cellular assays to profile and characterise Aurora A inhibitors has indicated an order of magnitude lower specificity^[18,31]^. A modest number of early studies have pursued orthogonal approaches to Aurora A inhibition not dependent directly on competition with ATP. Aurora A interaction with N-Myc (**Fig. 1A**) has been disrupted allosterically by ATP-competitive inhibitors and orthosteric competitors have identified for PPI site with functional binding partner proteins, such as TPX2 (**Fig. 1A**)^[33–37]^. It is established that kinase inhibitors that target sites other than the ATP-pocket can lead to improved selectivity and novel pharmacology^[38,39]^. Additionally, therapeutically targeted PPIs are less likely to accommodate mutations without loss of protein function, therefore reducing the potential for emergence of resistance^[40,41]^.

**Figure 1.**
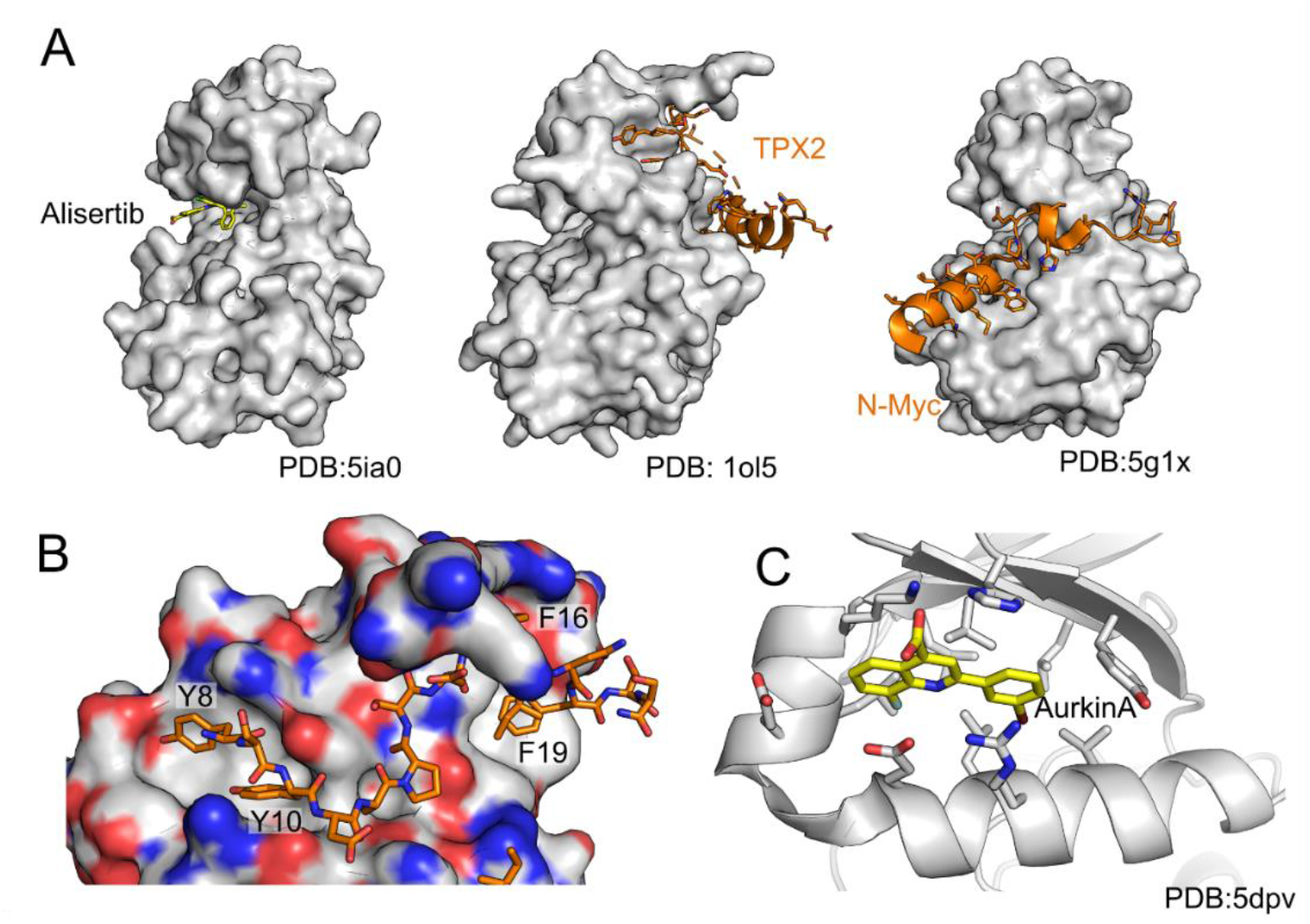
Aurora A interactions and inhibition. **(A)** Complexes of Aurora A with different interacting molecules. From left to right, ATP competitive clinical stage inhibitor alisertib (yellow carbons, PDB:5ia0^[43]^), TPX2 peptide (orange carbons, PDB:1ol5^[24]^and N-Myc (orange carbons, PDB:5g1x^[44]^) **(B)** Interaction of the N-terminal part of TPX2 peptide with Aurora A, with key aromatic residues labelled in the two pockets on the N-lobe. **(C)** Complex of Aurora A with Aurkin A (yellow carbons, 5dpv^[37]^), a low-affinity inhibitor of Aurora:TPX2 interaction.

Although ATP-binding site inhibitors that allosterically disrupt the interaction of Aurora A and N-MYC have demonstrated efficacy in xenografts^[42]^, to date, no reported PPI inhibitors of Aurora A-TPX2 have exhibited the potency or pharmacokinetics to be advanced to *in vivo* pre-clinical models. By targeting the TPX2 binding site unique to Aurora A, we aim to develop a small molecule inhibitor of Aurora A which is expected to show the therapeutic potential demonstrated by clinical agents such as alisertib and which additionally avoids the selectivity issues that typify ATP-competitive molecules. Moreover, by disrupting binding to a scaffolding protein TPX2, we hope to achieve also greater efficacy or new biological effects through the different mechanism of action.

## Results

### Development of Aurora A:TPX2 interaction inhibitors

#### Fragment-based drug design

We have pursued a structure-guided fragment-based drug development approach to develop inhibitors of the Aurora A:TPX2 interaction. Previous work from us and others have shown that the key interactions between Aurora A and TPX2 involve residues in the N-terminal half of the TPX2 epitope with mutation of tyrosines 8 or 10 or phenylalanine 19 resulting in significant drop in affinity for Aurora A^[34]^ (**Fig. 1B**). Also, the previously described Aurora A:TPX2 inhibitor Aurkin A binds to the so-called Tyr pocket inhibiting this interaction (**Fig. 1C**). This region of the TPX2 interaction does not overlap with where N-Myc binds to Aurora A.

Our aim was to develop potent inhibitors binding at this Tyr pocket, with properties that would enable *in vivo* evaluation of this approach to Aurora A inhibition. We started this process by screening a library of 600 fragments by thermal shift in the presence of an ATP-site binding inhibitor to focus fragment binding to sites other than ATP site. Thermal shift hits were progressed into a ligand-based NMR experiment, and a number of these such as 3-hydroxybenzoic acid (**1**) were shown to bind Aurora A and could be displaced by a TPX2 peptide fragment (amino acids 7-22) but not by a tight-binding ATP-site ligand. We established a competitive fluorescence polarisation (FP) assay with a longer, fluorescently labelled TPX2 peptide to mimic native-like interaction, but these NMR hits had no measurable activity in this assay. Moreover, we could not observe electron density for the fragments in X-ray crystallographic soaks. A focussed iteration of chemical elaboration of these hits yielded further fragments that maintained the desired competition profile in ligand-based NMR experiments, possessed activity in the FP assay, showing K_D_ values of around 1 mM and were confirmed to bind to Aurora A by isothermal titration calorimetry (ITC). Crucially, we were also able to obtain crystal structures of some of these hits in complex with the Aurora A protein, enabling structure-based drug design. A representative such fragment is compound **2**, a biphenyl molecule bearing a carboxylic acid and phenol group on one ring and a lipophilic trifluoromethoxy on the other. Compound **2** has a K_D_ 63 µM as measured by our competitive FP assay and K_D_ of 145 µM as determined by ITC. The binding of **2** to Aurora A, as determined by X-ray crystallography, alongside some key structural motifs showing both the ATP site and TPX2 peptide binding sites, is highlighted in **Fig. 2**. Our NMR and FP studies showed that these fragments are competitive with the TPX2 peptide (data not shown) and X-ray crystallography revealed that the hit fragments bind to part of the TPX2 binding site (**Fig. 2**), normally occupied by the Tyr8 and Tyr10 of TPX2 (we will refer to this pocket as the “tyrosine pocket” in the subsequent discussions).

**Figure 2.**
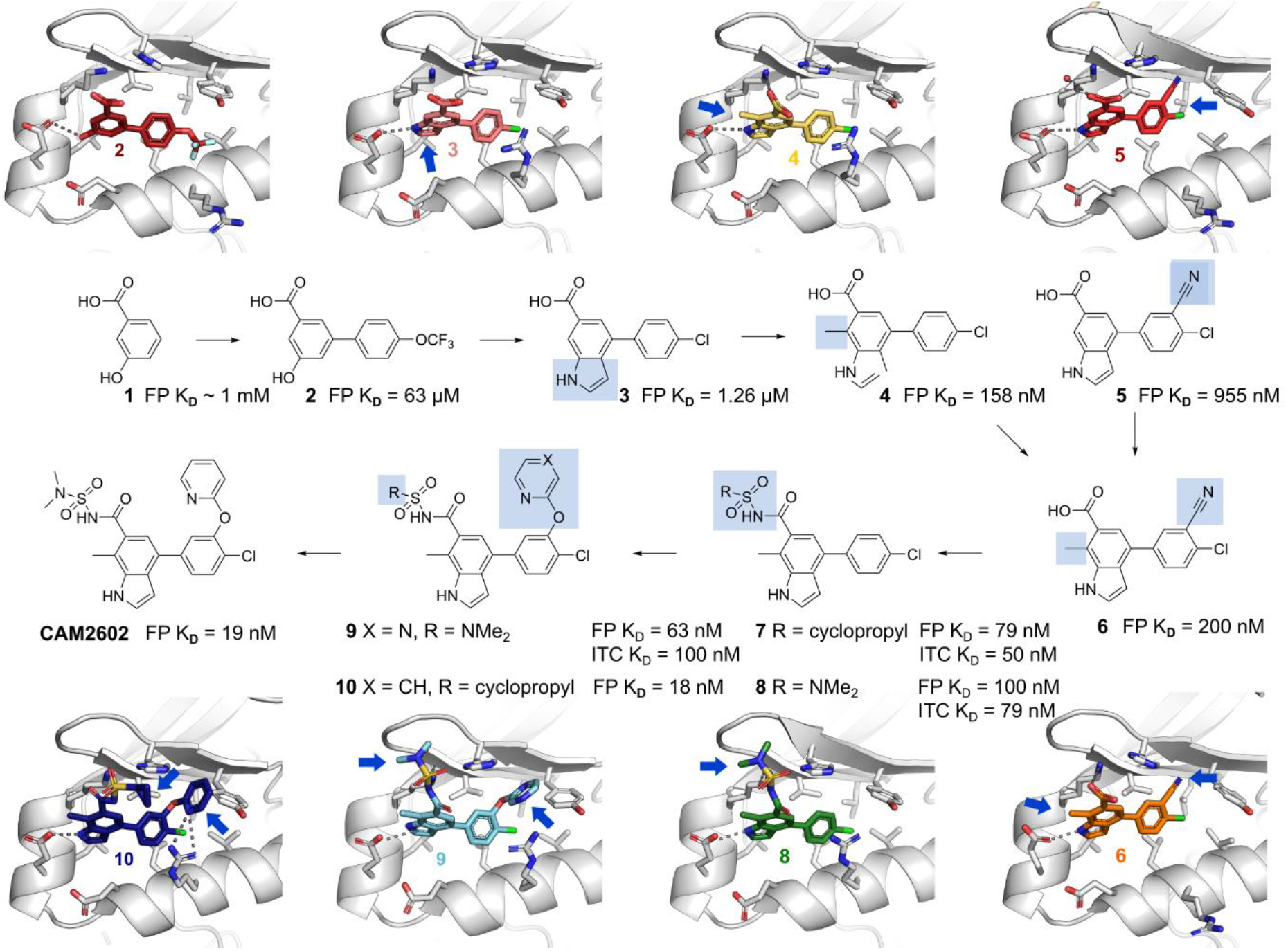
Aurora A:TPX2 interaction inhibitor design. Overview of the fragment-based development of **CAM2602** to inhibit the Aurora A:TPX2 protein-protein interaction. The blue boxes highlight the key change(s) at each step. Crystal structures of the compounds **2, 3, 4, 5, 6, 8, 9, and 10** in complex with Aurora A are show next to the chemical structures, with the key changes indicated by the blue arrows. The “meta-tunnel” is marked in the structure of **5**. The K_D_ values are obtained from a competitive FP assay or a direct ITC measurement, as indicated.

Through a further iterative development of the inhibitors utilising X-ray structure-based drug design and biophysics (FP and ITC), we improved the affinity of our weak, millimolar fragments hits by over 10,000-fold to generate the lead compound **CAM2602**. An early modification was to change the phenol group of **2** into indole whilst replacing the trifluoromethoxy with a smaller chlorine to give **3**, which improved the K_D_ to 1.26 µM (**Fig. 2**). The indole-aryl core of the molecule lays in a hydrophobic pocket assembled from Leu169, Leu178, Val182, Val206 and the side chain of Lys166. The indole nitrogen proton seems to form a hydrogen bond with the side chain of Glu170 thus mimicking the phenol of Tyr8 of TPX2. The carboxylic acid group was observed to interact with Lys166 and His201. Further, the electron density supported it being twisted from the plane of the indole ring in order to form a salt-bridge with Aurora A (**Fig. 2**). Our analysis of ligands in PDB and CSD^[45]^ databases show that carboxylic acids are more commonly in-plane with the aromatic ring (data not shown) and presumably this twisting incurs an energetic penalty upon binding. To minimise the loss of binding energy and to stabilise the torsional twist in the ground state, we introduced an *ortho* methyl group to **4**, which improved the K_D_ to 630 nM. We found that introduction of a *meta* nitrile group in the *para*-chloro ring led to a further modest improvement in potency and the crystal structure of Aurora A in complex with **5** revealed that the induced movement of Tyr199 generated a small pocket between Tyr199 and His201 (the “*meta*-channel”). Combining the modifications in **4** and **5** to give **6** resulted in reasonable FP activity and good cell permeability, permitting us to use **6** as a tool compound, particularly for early cell-based experiments. However, the potential utility of **6** *in vivo* is primarily hampered by poor hepatocyte stability, which was improved significantly through the introduction of isosteric replacements for the carboxylic acid, particularly acyl sulfonamides, in compounds **7** and **8**. In addition it was found that the *meta-*channel between Tyr199 and His201 could be further exploited by the replacement of the nitrile with an heteroaryl ether, to give **10** and lead compound **CAM2602** (**Fig. 2**,**3). CAM2602** engages with the tyrosine pocket through hydrophobic interactions at the bottom of the pocket and with polar interactions further outside. The indole NH hydrogen bonds with Glu170 side chain and the acyl sulfonamide stacks against His201 and Lys166. The pyridine ring in the meta position pushes Tyr199 sideways, creating a channel between His201 and Tyr199 and forming a T-stacked aromatic interaction with the latter. Finally, Arg179 latches on to the central aromatic ring, with **CAM2602** bound to a well-defined pocket which is partly induced by the binding of the inhibitor.

Our lead series maintains the acidic group present in fragment **2**, either as a carboxylic acid or an acylsulfonamide, whilst the phenol has been replaced with an NH in the form of an indole. An overlay of the crystal structures of the early hit **2** with **CAM2602** bound to Aurora A reveals a remarkable overlap of the core biaryl scaffold in the two compounds (**Fig. 3C**).

**Figure 3.**
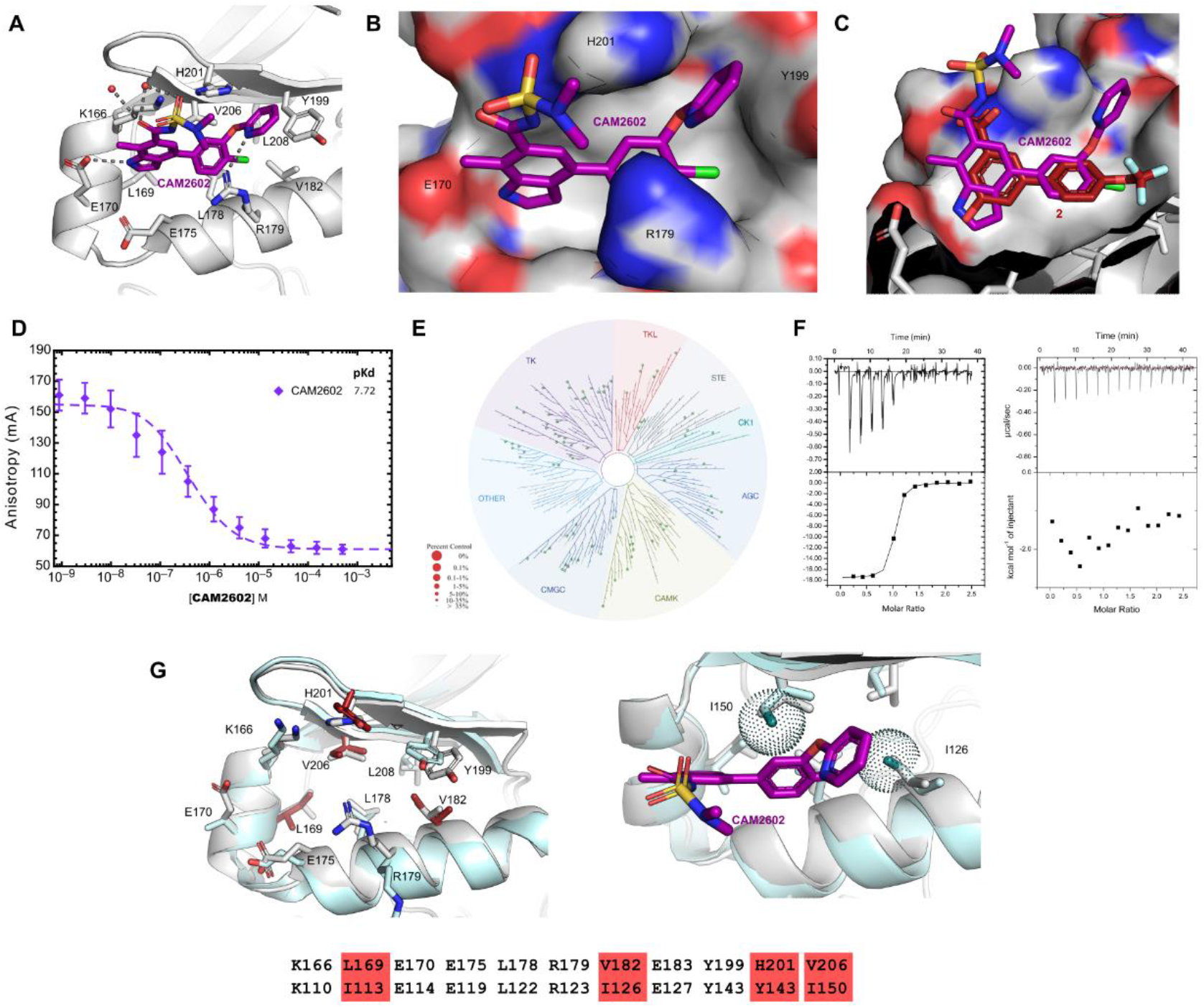
CAM2602 characterisation. **(A)** The X-ray crystal structure of **CAM2602** bound to Aurora A is shown (purple carbons; PDB: 8C1K) along with key interactions in the Tyr-pocket **(B)** View from above of the tyrosine pocket with Aurora A as a molecular surface. **(C)** Overlay of **3** and **CAM2602** showing remarkable preservation of the binding pose across the inhibitor development series. **(D)** Competition fluorescence polarisation assay of **CAM2602** competing with TPX2 peptide. **(E)** Kinase panel results using **9**. Red spheres would indicate cross-reactivity with kinases in the phylogenetic tree, with no observed reactivity for **9** in the panel of 97 human kinases. Details in **Figure S3 (F)** Isothermal titration calorimetry titration of **8** to Aurora A (left) and to Aurora B (right) **(G)** Conservation of residues in the Tyr pocket between Aurora A (light gray, PDB: 8C1K) and Aurora B (pale blue, PDB: 4AF3) with residues lining the tyrosine pocket shown as sticks and non-conserved residues in Aurora B coloured red. The same residues are shown below the figure with red background for non-conserved ones. On the left is a zoomed-in view of **CAM2602** binding to Aurora A, overlayed with the Aurora B structure (pale blue). The surface of additional methyl groups in Ile126 and Ile150 are displayed with surface dots, showing their proximity to **CAM2602**.

**CAM2602** displaces TPX2 from Aurora A with a <u>K_D_ of 19 nM and ligand efficiency of 0.33 (Fig 3D)</u>.

#### Kinase selectivity

We thoroughly evaluated the selectivity of our Aurora A:TPX2 inhibitors early on in the programme. Firstly, we tested **9** against 97 protein kinases in the DiscoverX KINOMEscan and failed to observe any detectable activity against these kinases at 10 µM, as expected from a non-ATP-site inhibitor (**Fig. 3C**).

Given our inhibitors bound to a PPI site, we hypothesised that they would show high selectivity for Aurora A over other kinases including Aurora B. Achieving selectivity over Aurora B has been recognized as a desirable feature of new drugs, but has thus far been challenging to achieve, due to the high sequence similarity (>70% identity) between the two kinase domains^[2,24,46,47]^ and the presence of a site that is analogous to the TPX2 binding site that, in the case of Aurora B, binds to the protein INCENP. To ensure our molecules did not bind to Aurora B, we measured binding of lead series representatives **7, 8** and **9** to both Aurora A and B by ITC. As expected, a good correlation is observed between the K_D_ of our inhibitors for Aurora A measured by competitive FP experiments and that from direct binding to Aurora A by ITC. Additionally, we observe an approximate 300-fold selectivity for Aurora A over Aurora B for **7** and **8** (**Fig. S2**). With the introduction of a meta-ether substituent in **9**, the compound’s potency against Aurora B was too weak to be measured – indicating greater than 1000-fold selectivity for Aurora A (**Fig. 3F**). The specificity of **9** for Aurora A over Aurora B is at least as great as the best compounds reported previously^[18,48]^.

The determinants of Aurora A vs B selectivity could be rationalised from our crystallographic data. Although many key residues that interact with their respective ligands are conserved, the shape of the base of the pocket is altered by three changes. In particular, His201, which in Aurora A is an important sidechain that forms a π-stack with the heterocyclic ethers and potentially participates in a charged interaction with the sulfonamide moiety in our lead compounds, is a tyrosine residue in Aurora B (Tyr145). Val182 and Val206 of Aurora A are both replaced by isoleucines in Aurora B, with the extra methyl groups making the Aurora B pocket somewhat smaller (**Fig. 3G**).

Potential toxicity of **CAM2602** was evaluated in protein-based Cerep panels, cellular toxicity assays, and peripheral blood mononuclear cells (PBMC) assays. High content cell toxicology of **7**, up to 40 µM in HepG2 cells, indicates that there were no measurable effects on cell growth, nuclear size, DNA structure, cell membrane permeability, mitochondrial mass, mitochondrial membrane potential or cytochrome c release (**Table S1**). Lead compound **CAM2602** exhibits only one off-target activity in the Cerep screen inhibiting binding of an agonist radioligand to human adenosine 3 (A3) GPCR by 55% at 10 µM. **CAM2062** does not inhibit hERG, or a panel of CypP450 enzymes at 25 µM (**Table S2**). Some of the ADMET properties of **CAM2602** are shown in **Table S3**.

#### Mechanistic characterisation of the Aurora A:TPX2 inhibitors

##### Target engagement in cells induces Aurora A mislocalisation

Previous reports have shown that Aurora A is recruited to the mitotic spindle through its protein-protein interaction with TPX2^[21,22]^. We have previously reported a high-content screening assay in which we can detect the displacement of Aurora A from the spindle in mitotic cells^[37]^. Here we used this assay to provide a measure of cellular target-engagement for our key compounds (**Fig. 4**). In parallel, we performed a related high-content assay measuring loss of the activating phosphorylation at threonine 288 (P-Thr288) on Aurora A. In agreement with previous data^[37]^, the EC_50_ values in these two assays were well-correlated (**Fig. 4A, B**).

**Figure 4.**
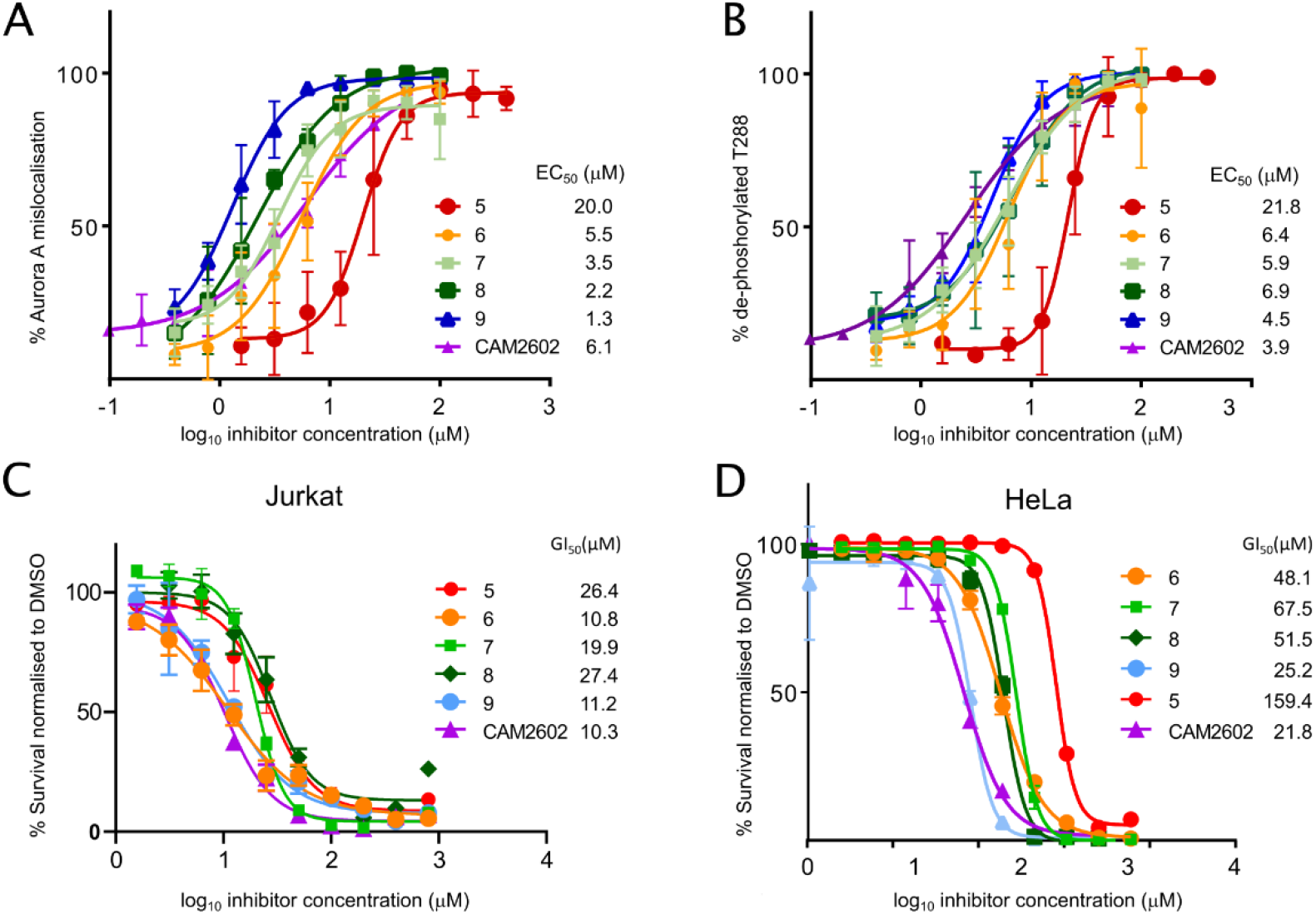
Cellular efficacy of the CAM2602 series. **(A)** High content microscopy assays to evaluate mis-localisation of Aurora A from the mitotic spindle or loss of phospho-Thr288 Aurora A in mitotic nuclei when treated with inhibitor. HeLa cells were treated with titrations of the indicated compounds for 2 hours before being fixed, stained for Aurora A and analysed using high-content microscopy to determine the percentage of observed mitotic cells at each concentration with spindle-displaced Aurora A (mislocalisation). The indicated EC_50_ values for each compound were calculated from the plots of assay scores against compound concentration. **(B)** As in A but stained for dephosphorylated Thr288 Aurora A. **(C)** Viability assays in Jurkat cells. Jurkat cells were cultured for 72 hours with titrations of the indicated compounds. Viability assays were performed following the treatment period and the data normalised to the vehicle-treated controls. GI_50_ values were calculated from plots of the viability assay data. **(D)** Viability in HeLa cells, determined similarly to (C).

An acute cellular consequence of inhibiting the mitotic function of Aurora A is the appearance of spindle abnormalities in those cells undergoing mitotic division^[49,50]^. Driven by deregulation of centrosome maturation and spindle pole forces, the abnormalities can be broadly characterised as including loss of spindle bipolarity and/or misalignment of the condensed chromosomes at the metaphase spindle; observations of these phenotypes have been used in pre-clinical and clinical studies employing ATP-competitive Aurora A inhibitors^[32,51,52]^. Treatment of HeLa cells with **6** for 6 hours resulted in significant increase in misaligned or trailing chromosomes, based on immunofluorescence microscopic analysis of chromatin DNA, Aurora A and α-tubulin (**Fig. S4A, B**).

#### Impact on viability in dividing cancer cells

Blocking the protein-protein interaction between Aurora A and TPX2 is predicted to disrupt Aurora A function in dividing cells^[20]^ leading to defects in spindle assembly, transient activation of the spindle assembly checkpoint and eventual apoptosis in a post-mitotic G1 arrest^[53]^. Actively cycling cells experiencing Aurora A inhibition are, therefore, expected to exhibit eventual loss of viability due to prolonged disruption of Aurora A function. The compounds were titrated in the growth assay to estimate their cytotoxic impact against either Jurkat acute T cell leukaemia cells or HeLa cervical adenocarcinoma. In general, we observed lower GI_50_s in compound treatments with Jurkat cells (**Fig. 4C, D**). To explore the potential therapeutic window for our compounds in dividing cancer cells versus normal tissues we made use of peripheral blood mononuclear cells (PBMCs). PBMCs are viable in tissue culture conditions, but do not cycle in the absence of a lymphocytic stimulus such as anti-CD3/CD28^[54,55]^. Non-cycling cells should not require active Aurora A, so assessing cell viability in the PBMCs may serve an indirect measure of potential off-target toxicity. We observed that most of the compounds with cell activity in HeLa and Jurkat cell viability experiments had no impact on the non-cycling PBMC cells when applied at less than 200 µM, which was an order of magnitude greater than the typical GI_50_ values seen in the equivalent Jurkat cell data (**Fig. S5**). As a control, the PBMC cells were also treated with ATP-competitive Aurora A inhibitor, alisertib, which also demonstrated no toxicity in the PBMC cells. Treatment with staurosporin, a non-selective kinase inhibitor that exhibits promiscuous cytotoxicity, resulted in dose-related killing PBMCs, confirming that the assay was capable of reporting non-specific cell-killing effects.

#### Biomarkers of Aurora A-TPX2 disruption

Phosphorylation of serine 10 on histone H3 (PH3) has been used as an indicator of mechanistic target engagement for ATP-competitive Aurora A inhibitor alisertib^[32,56–58]^. Aurora A inhibition produces a delayed G2/M transition driving accumulation of PH3 through the activity of Aurora B^[59,60]^. We treated Jurkat cells with either an early lead compound (**7**), alisertib or a vehicle control and followed PH3 levels over time by western blotting. Accumulation of PH3 in Jurkat lysates was observed from 16 hours following treatment both with alisertib and **7** (**Fig. S6A**).

It has previously been shown that PH3 accumulation in tumour cells treated with Aurora A inhibitors is detectable from as early as 4-6 hours with microscopy^[32,57]^. This suggests a sensitivity advantage for techniques that can resolve mitotic cells in asynchronous cell samples, so we next explored flow cytometry for detection of PH3 and phospho-Thr288 (P-T288) changes in Jurkat cells treated *in vitro* with varying GI_50_-multiples of **7** or a vehicle control for 8 hours. Supporting validation of PH3 immunostaining in these samples, this marker was only detectable in mitotic cells, identifiable by their 4n DNA. Samples treated with **7** demonstrated a consistent increase in PH3-positive mitotic cells compared to vehicle controls (**Fig. S7A, B**). A 2x GI_50_ dose of **7** yielded almost a 3-fold increase in mitotic cells compared to DMSO exposure, with a similar magnitude of increase at a 5x GI_50_ dose. Complementing the PH3 data, decreased P-Thr288 Aurora A was observed in the mitotic cells treated with **7**. This detection of biomarker modulation was repeated for the lead compound, **CAM2602**, with alisertib as a positive control using Jurkat cells *in vitro* (**Fig. 5A**). Under these conditions, both **CAM2602** and alisertib treatment exhibited similar evidence of inhibition of Aurora A phosphorylation.

**Figure 5.**
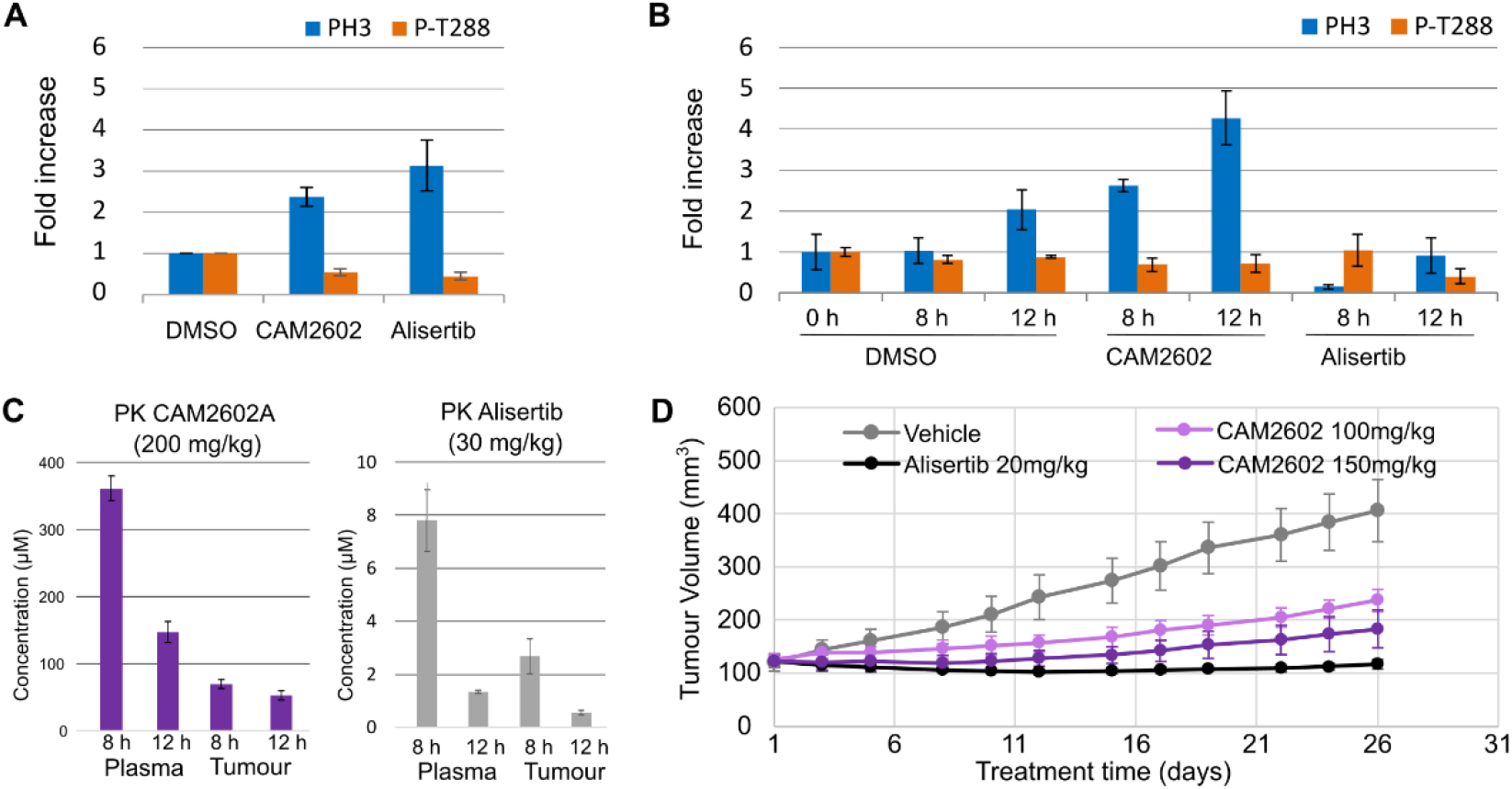
*In vitro* and *in vivo* characterisation of CAM2602. **(A)** Jurkat cells were treated for 8 hours with 20 µM **CAM2602** or 14 nM alisertib and analysed by flow cytometry for PH3 positive cells relative to vehicle controls. PH3-positive cells from each sample were assessed for loss of P-Thr288 positivity. **(B)** Female NOD scid gamma (NSG) mice bearing solid Jurkat tumours (subcutaneous implantation, rear dorsum) were administered a single oral dose of either **CAM2602** or vehicle. Tumour cells from 0, 8 or 12 hours of treatment were analysed by flow cytometry similarly to *in vitro* samples in panel A. **(C)** Pharmacokinetic analysis of CAM2602 or alisertib concentrations in tumour and plasma samples taken at 8 or 12 hours after dosing with 200 mg/kg and 30mg/kg, respectively. **(D)** NSG mice bearing subcutaneous, solid tumour xenografts of Jurkat cells were dosed orally once per day with either vehicle, **CAM2602** or alisertib, as indicated (n=5). Tumour volumes were estimated periodically over the 26 days of dosing by calliper measurement. Error bars show standard deviations from the mean.

#### PPI inhibitor of Aurora A-TPX2 demonstrates *in vivo* activity

Given the favourable ADMET profile of **CAM2602** (**Table S3**) and its ability to modulate biomarkers of target engagement *in vitro*, we next sought to demonstrate that **CAM2602** could affect tumour cell biomarker modulation *in vivo* following acute systemic administration in a mouse xenograft model.

We assessed first the pharmacokinetics of **CAM2602** by administering the compound at 3 separate doses in female CD-1 mice and measuring the total concentration of compound in plasma over time (**Fig. S8)**. The intravenous dose is cleared in a first-order elimination process. At higher doses, administered orally, the concentration of compounds rapidly reaches a plateau that is maintained for at least 8 hours. These clearance profiles suggest that one or more clearance mechanisms, i.e. efflux and/or metabolism may be saturated at these compound doses. The oral bioavailability of **CAM2602** at 50 mg/kg was 99.8% while no weight loss or adverse events were observed in any PK studies (**Fig. S9**).

For the xenograft model, Jurkat cells were engrafted as a subcutaneous, solid tumour in the flanks of NOD SCID gamma (NSG) mice. Xenografted mice were orally administered a single dose of 200 mg/kg **CAM2602**, 30 mg/kg alisertib or vehicle, based on our earlier PK data for **CAM2602** (**Fig. S8**) or previously reported studies using alisertib^[32,61,62]^. Tumour and plasma samples were then taken 8- or 12-hours post-dosing. Resected tumours were digested into single cell aspirates, fixed and processed using flow cytometry to detect modulation of PH3 and P-Thr288 biomarkers (**Fig. 5B**). At both 8- and 12-hours post-dosing, xenografted tumour cells from **CAM2602**-treated mice demonstrated fold-increases in PH3 over vehicle controls matching those seen previously *in vitro* (**Fig. 5A, B**). Across the **CAM2602**-treated tumour samples, decreases in the Aurora A P-Thr288 marker were also evident, but changes to this marker were considerably less pronounced than those seen for *in vitro* conditions and were not significant. Plasma and tumour concentrations of **CAM2602** exhibited high micromolar concentrations of the compound in both compartments at both 8- and 12-hour time points (**Fig. 5C**). When adjusted for mouse plasma protein binding (**Table S3**) the predicted free drug concentrations in plasma (5.4 µM at 8 hours and 2.2 µM at 12 hours) are well in excess of the K_D_ (20 nM) for the target, supportive of likely target engagement. Moreover, the measured tumour concentrations (70 µM at 8 hours and 54 µM at 12 hours) suggest meaningful tissue exposure consistent with levels required for inhibition in cells up to 12-hours post-dosing. Contrary to our *in vitro* data (**Fig. 5A**), tumour samples recovered from alisertib-treated mice yielded a decrease in PH3 at 8 hours, and neither 8- or 12-hour samples yielded the increase in PH3 expected from Aurora A inhibition (**Fig. 5B**). Tumour and plasma PK measurements 8- and 12-hours post dosing with alisertib indicated either micromolar or very high nanomolar tissue concentrations for this potent inhibitor (**Fig. 5C**). Alisertib is likely to have off-target activity against Aurora B at these high concentrations, which might be expected to decrease PH3, therefore overriding the increase in PH3 expected from Aurora A inhibiton^[60,62]^.

#### CAM2602 induces growth suppression of tumour xenografts

Tolerability studies with 50, 100 and 150 mg/kg administered to NSG mice (daily dosing for 7 days, followed by 7 days without dosing) indicated that the highest dose examined of 150 mg/kg was tolerated without overt toxicity (**Fig. S9**). An efficacy study was performed using xenografted NSG mice bearing subcutaneous Jurkat cells implanted as solid tumours with a daily oral dose of either 100 or 150 mg/kg **CAM2602**, 20 mg/kg alisertib or vehicle for 26 days. Tumour volume measurements were taken three times per week during this time. The volume data indicated that vehicle-treated mice exhibited continuous tumour growth during the study, whereas the two doses of **CAM2602** were capable of successfully reducing tumour growth, the higher of the two doses having the greater effect (**Fig. 5D**). Alisertib had the greatest impact on tumour growth, likely due to the higher potency of this inhibitor. In agreement with earlier assessments of toxicity, there were no observations of toxic phenomena among the treated mice for the duration of the study and no evidence of loss of body weight (data not shown). Inhibition of Aurora kinases with ATP-competitive inhibitors has previously been linked to dose limiting toxicities such as bone marrow ablation and neutropenia^[17,47]^. Possible loss of blood cell lineages indicative of such toxicities were additionally analysed using blood samples taken from all mice upon completion of the efficacy study. These analyses indicated a very mild anaemic response in all non-vehicle animal dosing groups with a slight drop in haematocrit readings, but this was coincident in all cases with an elevation in reticulocyte count (**Fig. S10)**.

Aurora A overexpression is known to drive resistance to taxanes in cancer cells^[12,13,63]^. In addition, compelling data indicates that inhibition of Aurora A synergises with paclitaxel in cell lines exhibiting Aurora A amplification^[64]^. Using an earlier compound in our series, **6**, with an analogous structure and mode of action to **CAM2602**, we were able to demonstrate drug synergy with Taxol in the pancreatic cell line PANC-1, emulating benefits previously observed for ATP-competitive Aurora A inhibitors (**Fig. S11**). Considering the greatly limiting toxicities associated with Taxol in the clinic, a major therapeutic implication of these results could be the potential to greatly reduce required doses of Taxol when applied in combination with a drug targeting the Aurora A-TPX2 PPI. A prediction for Aurora A inhibition, including PPI-targeting agents, is the reversal of taxane resistance, which suggests a promising clinical opportunity to treat tumours with combinations of these agents^[12,13,63,64]^. Taxane resistance is a major clinical challenge with nearly half of all patients exhibiting primary resistance or eventually relapsing with treatment-resistant disease; agents that reverse taxane resistance would find utility in epithelial ovarian cancers, mammary adenocarcinomas and non-small cell lung carcinomas, for example^[65–68]^.

## Discussion

Small molecule inhibition of Aurora A is an attractive strategy for the treatment of a wide range of human malignancies^[3–5,12,14–16]^. Consequently, several high-potency, orthosteric, ATP-competitive inhibitors of Aurora A have been developed^[17]^. Encouraging trial data have been seen for one such inhibitor, alisertib, across a range of cancers, but significant dose-limiting toxicities are consistently observed^[31]^. The promise of PPI inhibitors of kinases is that they bind to less conserved sites in the target and are more likely to exhibit better selectivity than orthosteric ATP-competitive molecules^[38,69]^. Therefore small molecule inhibitors targeting PPIs potentially exhibit fewer off-target toxicities and can have reduced propensity to develop resistance in cancer cells^[38–40]^. TPX2 is a particularly promising binding partner to block in this way, exhibiting a broad repertoire of activity-promoting properties in relation to Aurora A^[1,20,24]^.

We have developed through fragment-based, structure-guided approaches a series of novel compounds that inhibit the PPI between Aurora A and TPX2. The initial fragment hits identified from screening with the ATP site blocked by a high affinity inhibitor, were very weakly active, but, guided by continuous crystallographic analysis of the inhibitors in complex with Aurora A, we were able to increase target affinity by more than 10,00-fold, clearly demonstrating the ability of fragment-based and structural biology approaches to develop potent PPI inhibitors when a suitable binding pocket is present. These compounds occupy a hydrophobic pocket on the surface of Aurora A, discrete from its ATP-binding catalytic site, which forms the interaction surface for a linear N-terminal segment of the interacting peptide from TPX2. They displace critical interactions made by the Tyr8 and Tyr10 residues of TPX2 with Aurora A, directly inhibiting the binding of TPX2 to a key hotspot in Aurora A^[34,70]^. Notably, the compounds interact with Aurora residues that are not conserved in the closely related Aurora B kinase, providing a structural rationale for their high selectivity.

These are the first high-affinity ligands inhibiting this allosteric site and our lead compound **CAM2602** has pharmacological properties that enable it to be used in *in vivo* studies. We find that these compounds are cytotoxic to cancer cells alone or in a synergistic combination with paclitaxel, with their cytotoxic effects proportional to target engagement marked by Aurora A mislocalisation and dephosphorylation on Thr288.

In a solid tumour xenograft model oral delivery of **CAM2602** successfully elicited biomarkers of target engagement, increasing PH3 positive cells and decreasing the proportion of those cells positive for P-Thr288 Aurora A, moreover this compound also reduced tumour growth. These results show that an inhibitor of the Aurora A-TPX2 PPI is a viable route to therapeutic intervention in cancer.

The lack of overt toxicity seen *in vitro* and particularly in the *in vivo* studies with lead compound **CAM2602** is noteworthy. Considering the high doses required to deliver our target tumour drug levels, it was possible that toxicity similar to that seen with ATP-competitive Aurora A inhibitors in the clinic^31^ might impact the practical utility of **CAM2602** in the sustained multi-dose efficacy study. This apparent lack of toxicity may reflect the particularly high target-specificity which is characteristic of enzyme inhibition by the PPI mode rather than at the ATP-binding pocket^[38,39]^.

In conclusion, we have developed a small molecule inhibitor of the Aurora A:TPX2 interaction, for which we provide a first example of efficacy in a xenograft model, providing a proof of concept for further development. In addition, the encouraging *in vitro* synergy demonstrated with Taxol suggests an important clinical modality for this new class of inhibitors.

During the course of this work, Bayliss and co-workers have published the results of two crystallographic fragment screens against Aurora A^[34,35]^. Our target pocket, where tyrosines 8 and 10 of TPX2 bind, was identified as one of the hot spots for this PPI and a number of diverse fragments were found in this pocket, providing possibilities for further development of Aurora A:TPX2 inhibitors.

## Abbreviations

SAR: Structure Activity Relationship;

## Acknowledgements

Chris Abell led this project throughout, but passed away in 2020 while the manuscript was in preparation.

We’d like to thank Dr George Trainor, our drug discovery advisor for the SDDI award, as well as Dr Philip Jordan and Prof. Steve Wedge for helpful discussions and advice.

We are grateful for Diamond Light Source for access to beamlines I04-1, I03 and I24 (proposals mx9537 and mx14043) ESRF for access to beamlines MASSIF-3 and ID29 and Synchrotron Soleil for access to Proxima-2 beamline, data from which contributed to these findings. We thank Biophysical and X-ray crystallographic facilities at the Department of Biochemistry for access to instrumentation and technical support.

## Funding

This work was funded by a Wellcome Trust Strategic award (090340/Z/09/Z) and a Wellcome Trust Seeding Drug Discovery Initiative award (101134/Z/13/Z).

## Author Contributions

JS, CA, DS, MH, TLB and ARV – envisaged the project, wrote the grant application and supervised the work. JS and DES led the project at different stages. DES, TPCR, JF, RS, CD, EA, and JS contributed to compound design and/or chemical synthesis. AH was in charge of modelling and data management and contributed to compound design. GF, MR, MR and MH were in charge of crystallography and 3D structure interpretation. TM, MR, BB and XW were responsible for protein biochemistry and biophysical analyses. SRS, EGA, AA-L and GM were responsible for cell biology and animal experiments. TP advised on medicinal chemistry. SRS, DES, JS, MH and ARV wrote the paper. All authors have edited the manuscript and contributed to their part of the data analyses.

## Notes

### Competing Interest Statement

The authors have declared no competing interest.

### Summary of Updates

We have revised the manuscript after rejected submission to a journal, taking referees points into account and focusing the paper more on the development of inhibitors and moving some of the biological validation of intermediate compounds to the supplement. At the same time we changed corresponding authors to reflect the change of manuscript focus.

