## Supplemental data for "Selective Aurora A-TPX2 interaction inhibitors have *in vivo* efficacy as targeted anti-mitotic agents"

### Selective inhibitors of the Aurora A-TPX2 protein-protein interaction exhibit in vivo efficacy as targeted anti-mitotic agents

Supplementary material

#### Contents

Methods ... 3

Cell culture ... 3

Viability assays ... 3

High-content screening ... 3

Confocal microscopy ... 4

Flow cytometry ... 4

Western blotting ... 5

*In vivo* studies ... 5

Synergy analysis ... 6

Protein expression ... 6

Fluorescence polarisation (FP) assay ... 7

Isothermal titration calorimetry ... 7

Crystallisation and structure determination ... 7

Biophysical, cellular and in vivo analyses ... 8

Figure S1. Fluorescence polarisation anisotropy assay. ... 8

Figure S2. ITC analysis of selectivity for Aurora A vs. Aurora B for compounds 7,8, and 9. ... 9

Figure S3. DiscoverX KINOMEscan specificity screen of compound 9. ... 10

Table S1. High content toxicology analysis. ... 11

Table S2. Cerep Express Profile screen data. ... 11

Table S3. Calculated and measured ADMET properties for CAM2602 ... 13

Figure S4. Mitotic spindle abnormalities in cells treated with 6. ... 14

Figure S5. Toxicity in non-cycling cells. ... 14

Figure S6. PH3 levels evaluation with compound 7. ... 15

Figure S7. Flow cytometry gating strategy to detect Aurora A inhibition biomarkers. ... 15

Figure S8. Pharmacokinetics of CAM2602. ... 16

Figure S9. Tolerability of CAM2602. ... 16

Figure S10. Analysis of blood samples upon conclusion of efficacy study. ... 18

Figure S11. Aurora A:TPX2 PPI inhibitors synergise with Taxol in PANC-1 cells. ... 19

Table S4. Crystallographic data collection and refinement parameters. ... 20

Figure S12. Electron densities of ligands ... 24

Figure S13. High-content assay to assess cellular Aurora A engagement by PPI inhibitors. ... 25

#### Methods

##### Cell culture

HeLa, PANC-1 and Jurkat cells were maintained in humidified incubators at 37 °C, 5% $CO_2$ using either DMEM (HeLa and PANC-1: high glucose, GlutaMAX™ Supplement, pyruvate; ThermoFisher Scientific 10569010) or RPMI 1640 (Jurkat and PBMC: GlutaMAX™ Supplement, HEPES; ThermoFisher Scientific 72400021) media supplemented with 10% foetal bovine serum. As a positive control in the high-content screening assays we made use of a previously reported stable HeLa FlpIn TREx cell line expressing a fusion mCherry-TPX2-1–43 protein which was inducible upon addition of doxycycline (0.5 mg $ml^{-1}$)[37]. New vials of PBMC cells were obtained for each viability experiment (ATCC, PCS-800-011).

##### Viability assays

Cells were seeded onto sterile, flat-bottomed, 96-well tissue culture plates in antibiotic-free media; HeLa were seeded the day before treatment at a density of $5x10^3$ per well, whereas Jurkat and PBMC cells were seeded at $2x10^4$ or $1x10^5$ per well, respectively, on the day of treatment. All wells per plate contained 100 µl of cells and/or media and the outermost wells of each plate contained media-only controls. On the day of treatment, 10-point, 2-fold dilution series of each compound were prepared in antibiotic-free media on separate, sterile, round bottomed 96-well plates. All series concentrations were adjusted to 5-fold higher than the intended final concentrations before 25 µl of these were then pipetted in triplicate to the flat-bottomed plates with cells, yielding a final volume of 125 µl per well. Matching DMSO-treatment dilution series were included in triplicate on each plate. Media-only edge wells received 25 µl of media to maintain equal final volumes across all wells on the plates, which were then sealed with sterile, breathable membranes beneath the plate lids and incubated in humidified incubators at 37 °C, 5% $CO_2$ for 72 hours. Depending on cell line, cell growth per well was assessed using the CellTiter-Blue assay (Jurkat cells, Promega) or sulforhodamine B assay (Hela). Cell-free control wells were used to calculate assay blanks for subtraction from assay values per treatment condition per plate; triplicate means of corresponding DMSO control well assay values for were used to determine fold-survival values for each compound treatment condition. $GI_{50}$ values were calculated from four-parameter dose-response curves that were fitted using Prism GraphPad software (La Jolla, CA).

##### High-content screening

The high-content imaging Aurora A mislocalisation and Thr288 dephosphorylation assays have been described previously by our lab[37]. Briefly, 24 hours after seeding $8x10^3$ HeLa cells in 100 µl media per well of tissue-culture treated 96-well plates (ThermoFisher, 167008), the cells are treated with 9-point, 2-fold titrations of compound in media for 2 hours under standard tissue culture conditions. Drugging volumes were managed as described above for the viability assays (i.e. 25 µl is added to a final volume of 125 µl on cells to yield 5x dilution). Drugging media was supplemented to give a final concentration of 10 µM Velcade (Bortezomib, Selleck Chemicals) to reduce numbers of anaphase cells yielding false-positivity during image analysis. Following 2 hours incubation under drugging conditions, the plates were aspirated, fixed, permeabilised and stained as before.

Imaging of the plates was performed on an ImageXpress Micro Confocal High-Content Imaging System (Molecular Devices) using a 20x ELWD objective (optimal for 96-well plates with standard, 1.9 mm thick transparent bases) and laser autofocussing per field. For each well 12 non-overlapping fields in 3 fluorescent channels were acquired with bright-field optics and 2x2 binning, which allowed for approximately 100 mitotic cell observations per triplicate well. Custom Module Editor

(CME) image analysis software (Molecular Devices) was used to quantify mitotic cell phenotypic responses, which were used to calculate assay endpoints.

Aurora A mislocalisation assay image data was analysed in CME by using Hoechst/DAPI channel image data to locate all individual nuclei per field. Corresponding TPX2/CY5 channel image data were used to identify the mitotic cell sub-populations in each field through TPX2-positivity of their nuclei. Intensity thresholds >100 times that of the image background were set in CME to distinguish DAPI and FITC channel signal from any noise. For each mitotic nucleus a top-hat filter with a 25 µm kernel was used to define a fine mitotic spindle mask. Per mitotic spindle mask, the corresponding average Aurora A/FITC channel intensity was measured. The resulting cell-level data was exported and analysed in Excel whereby the highest spindle Aurora-A intensity in the darkest 10% of mitotic cells from untreated control wells was used to set a per-plate assay threshold below which Aurora-A was classified as delocalised from the spindle. The assay threshold was then applied across all mitotic cells recorded per well, the percentage of cells with Aurora-A intensity in the spindle mask below the threshold was reported as the percentage of mitotic cells per well with mislocalised Aurora A. The Thr288 dephosphorylation assay was performed and analysed the same way as for the mislocalisation assay, but substituted PH3 and P-Thr288 Aurora A antibodies for TPX2 and total Aurora A, respectively. In this case, PH3-positivity was used to identify mitotic cells and the mitotic spindle mask was replaced with a whole-nucleus mask for the purpose of measuring P-Thr288 loss. A percentage of mitotic cells per well exhibiting dephosphorylated Thr288 Aurora A measure used the same assay threshold calculation as used for the mislocalisation assay. Diagram of the imaging scheme and image analysis are shown in Supplemental **Fig. S11**.

**Confocal microscopy**
HeLa cells were grown on sterile type-I borosilicate glass cover slips placed in 6-well tissue culture plates with $2x10^5$ cells per well. 24 hours following seeding, the cells were treated as indicated, then the media was aspirated and the cells were fixed using ice-cold methanol for 10 minutes. Fixed cells were permeabilised with 0.1% Triton-X100, 0.1% TWEEN20 in PBS for 10 minutes at room temperature before being washed in blocking buffer (3% BSA, 0.1% TWEEN20 in PBS) for 30 minutes. Anti-Aurora A (Abcam, ab52973, 1:500) and anti-tubulin (Abcam, ab6160, 1:500) were diluted in blocking buffer and used to probe the cells for 30 minutes at room temperature. Excess antibody was washed with 3 rounds of 0.1% TWEEN20 in PBS, followed by probing with secondary antibodies (goat anti-rabbit Alexafluor 488, A11034, 1:500; goat anti-rat Alexafluor 647, A21247, 1:500, ThermoFisher Scientific) applied and washed as per the primary antibodies, supplemented with 4 µg/ml Hoeschst 33342. Imaging was performed on a Leica SP5 confocal microscope using a 100 × 1.4 NA oil objective. Maximum projection images were created with z-stacks taken at 1 µm intervals. Pixel intensities were kept sub-saturation. Laser exposure and detector settings were identical across an experiment to allow comparison between samples.

**Flow cytometry**
Jurkat cells from either tissue culture or resected tumour xenografts were washed, fixed and permeabilised using reagents from BD Biosciences (Stain Buffer, 554657; BD Cytofix, 554655; Perm Buffer III, 558050). Ideally, $1.5x10^6$ cells per sample were washed once with 500 µl cold Stain Buffer and transferred to clean 1.5ml centrifuge tubes. Samples were pelleted and aspirated before fixing with 250 µl BD Cytofix buffer following a brief vortex in the fixative and incubation on ice for 15 minutes. The fixed cells were then washed as before and subsequently pelleted and aspirated prior to being permeabilised by slow addition of 500 µl cold Perm Buffer III while vortexing. Samples were incubated on ice for 30 minutes then washed as before. The cells were then sequentially stained in three steps with anti-Aurora A P-Thr288 (1:100, Cell Signaling #3079), goat anti-rabbit Alexafluor555

(1:500, Life Technologies A21429) and finally AF647-conjugated anti-histone H3 (phospho-S10, 1:400, Cell Signaling #3458). For resected xenograft samples, AF488-conjugated human specific anti-CD3 (1:200, BD Pharmingen 557694) was included in the final staining step to allow exclusion of possible host cell contamination. The sequence of antibody staining is required to avoid species cross-reactivity between the chosen antibodies. The antibodies were applied to the cell samples in 100 µl of staining buffer, incubated for 30 minutes at room temperature with rotation and washed twice in 500 µl of stain buffer between each antibody step. Cells remained in the final wash supplemented with 4 µg/ml Hoechst 33342 and 250 µg/ml RNAse A. The cells were transferred to flow cytometry tubes and incubated in the dark at room temperature for 30 minutes before being analysed. Analysis of flow cytometry samples was performed on a BD LSRFortessa equipped to excite the samples at 355nm, 488nm and 640nm and to resolve the fluorescent probes using separate detectors. Experiment data was analysed using FlowJo Ver.10 software (FlowJo, LLC). Gating strategies are shown in **Fig. S6**.

**Western blotting**
Total protein was isolated by directly lysing the cells in non-denaturing lysis buffer (50 mM HEPES-HCl pH7.4, 250 mM NaCl, 0.2% Triton X-100, 1 mM EDTA, 1 mM dithiothreitol, 1 mM NaF, 10 mM β-glycerophosphate, 0.1 mM $Na_3VO_4$, 1x Roche cOmplete protease inhibitors). Protein lysates (12 µg per lane) were resolved on SDS-PAGE gels, transferred onto an Immobilon-P, PVDF membrane (0.45 µm, Millipore), and probed with either anti-histone H3 (1:1000, NEB, 9715S) or anti-histone H3 (phosphor S10, 1:2000, Abcam, ab14955). Secondary HRP-conjugated antibodies were used (GE Healthcare) and the signal was detected using an Amersham enhanced chemiluminescence system (ECL, GE Healthcare).

***In vivo* studies**
*In vivo* pharmacodynamics, tolerability and efficacy studies were carried out by Axis Bioservices Ltd (Northern Ireland). Pharmacokinetic work was done at WuXi AppTech (China). Female CD-1 mice were used in pharmacokinetics studies and female NOD-SCID gamma (NSG) mice were used for all other *in vivo* studies. For xenograft studies, Jurkat E6.1 cells (ATCC) were bulk-grown in RPMI 1640 media (GlutaMAX™ Supplement, HEPES; ThermoFisher Scientific 72400021) supplemented with 10% foetal bovine serum. Tumour cell implantation employed $2x10^7$ cells in matrigel per tumour, injected subcutaneously to the rear dorsum. Tumour volumes post-implantation were monitored using calliper measurements and mice were advanced for treatment when tumour volumes between 150mm-200mm$^3$ were reached. Where used, compounds were formulated in DMSO:20% HP-β-CD (2-hydroxypropyl-beta-cyclodextrin in PBS, 2.5:97.5) with pH adjusted to 7.6. All treatments were administered by oral gavage.

For pharmacodynamic biomarker studies, mice aged 5-7 weeks at time of implantation were administered single doses of the indicated treatments and were harvested for tumour resection and collection of whole blood by cardiac puncture at 0, 8 or 12 hours post-dosing. Plasma samples were submitted for PK analysis (Xenogesis Ltd.). Resected tumours were digested to single cell aspirates in dissociation buffer (RPMI medium supplemented with 5% FBS, Collagenase type I (200 U/ml) and DNase I (100 µg/ml)) for 30 minutes at 37°C with periodic vortexing and passed through a 70 µm filter with PBS washes. Tumour samples were cryogenically frozen and stored prior to being processed for flow cytometry as described above. Efficacy studies employed xenografted mice aged 6-8 weeks. Dosing was applied daily for 26 days and tumour volumes ($4/3\pi r^3$) were recorded three times per week by calliper measurements using three reference diameters to estimate geometric mean diameter. Samples were harvested 8 hours after the final dose. Tolerability studies used nonxenografted mice aged 6-8 weeks. Doses were applied daily for 7 days followed by a 7 day period with no treatment. Animal bodyweight, behaviour and appearance were monitored daily.

**Synergy analysis**

Drug synergy experiments using the Bliss independence model were performed as previously reported[64]. 96-well plates were seeded with $5x10^3$ PANC-1 cells per well 24 hours prior to drugging with a dilution series of each drug in an 8x8 checkerboard pattern of combinations. For both drugs, the lowest drug concentration value in each series was a no-drug vehicle control, which allowed for true single-agent dosing to be represented among the permutations of drug ratios tested. After SRB staining to obtain the growth inhibition data, we used SynergyFinder webserver (https://synergyfinder.org/)[71] to identify synergistic drug combinations. The single-agent inhibition values were used to calculate a drug combination surface under the assumption of an additive effect. Regions of synergy were then detected by comparing observed combination data with the corresponding predicted values assuming additivity. In the final synergy plots, positive values indicate synergy regions, whereas negative difference values identify antagonistic effects.

**Protein expression**

Aurora A was expressed from pBAT4 or pHAT4 plasmid[72] in double cistronic construct with λ phosphatase, without which Aurora A was toxic to *E. coli*. Aurora A for biophysical assays was expressed from plasmid pBAT4-AurAS.003 which encoded for the kinase domain only (residues 126-390) of human Aurora A (Uniprot: O14965) followed by hexa-His tag. Deletion of the N-terminal localization domain implied the additional benefit of removing a region of the protein that was predicted to be intrinsically disordered. Further tailoring of the construct N- and C-termini was based on expression levels. For crystallography Aurora A contained also mutations Thr287Ala or Cys290Ala to reduce heterogeneity by activation loop phosphorylation and disulfide bond formation, respectively. For earlier compounds, a longer (residues 126-391) version of the protein without a C-terminal His-tag was used for crystallisation, as described in Janecek et al.[37] Aurora B protein was expressed from plasmid pNIC28-AurB (Addgene 39119).

Aurora A and Aurora B proteins were prepared with the same protocol. The protein expression was carried in BL21(DE3) strain (which was supplemented with pUBS520 plasmid for rare-Arg codon compensation for Aurora A) in 2YT media with 100 µg/ml of ampicillin. The cells were grown in shaker flasks to OD of 0.8-1.0 and expression induced with 400 µM isopropyl-thio-β-glycopyranoside for 3 hours at 37 °C. Cells were harvested by centrifugation and pellets stored at -20 °C. Cells were resuspended in 50 mM HEPES pH 7.4, 1 M NaCl, 100 mM Mg Acetate, 1mM ATP/1mM ADP, 25 mM Imidazole, 5 mM β-mercaptoethanol, with one tablet of protease inhibitors (cOmplete Protease Inhibitor Cocktails, Roche) and 500 µl of 2mg/ml DNaseI (Sigma: DN25). Cells were lysed with sonication or using an Emulsiflex homogeniser and lysate clarified by centrifugation at 30,000 *g* for 30 mins at 4 °C. The supernatant was filtered and protein purified with automated two-step protocol using an ÄKTA Pure chromatography system. The protein was captured in 5 ml FF HisTrap column (Cytiva) and washed with 50 mM HEPES/Na pH 7.4, 500 mM NaCl, 100 mM magnesium acetate, 1 mM ATP/1 mM ADP, 40 mM Imidazole, 5mM β-mercaptoethanol, 10% v/v glycerol until baseline stabilised. Protein was eluted in reverse flow with 50 mM HEPES/Na pH 7.4, 500 mM NaCl, 100 mM Mg Acetate, 1 mM ATP/ 1 mM ADP, 600 mM Imidazole, 5 mM β-mercaptoethanol, 10% v/v glycerol and the eluted protein directed to injection loop and injected directly to HiLoad 16/60 Superdex 75 pg column (Cytiva) which had been equilibrated with 50 mM HEPES pH 7.4, 50 mM NaCl, 100 mM Mg Acetate, 1 mM ADP, 0.5 mM TCEP, 10% v/v glycerol and column ran at 1 ml/min. Peak fraction was pooled, concentrated and stored in flash-frozen aliquots at -80 °C.

TPX2 peptide (residues 7-43, Uniprot: Q9ULW0) with C-terminal GGGCSS tail was expressed in *E. coli* as a GB1 fusion with an N-terminal His-tag and HRV 3C protease cleavage site for tag removal in vector pOP3BP, as described above. A pellet from 2 litre culture was resuspended in 50 mM HEPES pH 7.4, 500 mM NaCl, 40 mM imidazole, 10% glycerol, 0.5 mM TCEP and 500 µl of DNaseI (2 mg/ml) and lysed using a sonicator. Lysate was centrifuged for 30 min at 15,0000 *g* and filtered supernatant loaded on 1 ml gravity flow Ni Sepharose column (Cube Biotech). After washing with lysis buffer, the protein was eluted with 50 mM HEPES pH 7.4, 500 mM NaCl, 300 mM imidazole, 10% glycerol, 0.5 mM TCEP. Peak fractions were pooled and buffer exchanged with PD-10 column to remove imidazole and glycerol. Alexa Fluor™ 488 C5 Maleimide (catalogue no. A10254, Thermo Fisher Scientific) was added to the protein sample in 25-fold molar excess to label the C-terminal cysteine for 2 h at room temperature. Reaction was terminated with excess cysteine and protein cleaved with HRV 3C protease overnight. The cleaved protein was passed through second Ni Sepharose column to remove fusion protein and uncleaved material. Labelled peptide was purified by reversed phase chromatography using HiChrom 300 Å 4.6x250 mm C18 column with gradient elution from 10 % acetonitrile, 0.1 % trifluoroacetic acid to 90 % acetonitrile at 3ml/min flow rate, dried under vacuum, resuspended in 50 mM HEPES pH 7.4, 100 mM Mg acetate, 50 mM NaCl and stored at -80 °C in dark.

**Fluorescence polarisation (FP) assay**

The FP assay was done using a BMG Pherastar FS plate reader with a gain of 20% and target 90 mP. The Kd for TPX2 binding to Aurora A was determined to be 1.2 nM by direct titration of up to 200 nM of Aurora A protein to 11 nM labelled TPX2 peptide in 100 mM HEPES pH 7.4, 100 mM magnesium acetate, 50 mM NaCl, 0.02% P20, 1 mM DTT, 1 mM ATP, 10% (v/v) DMSO. The competition FP assay was run in the same buffer with 10 nM TPX2 peptide and 30 nM Aurora A. 12 concentrations of compounds from 1 µM to 2 mM were used as competitors in triplicate. The data was monitored for both anisotropy and for change in total fluorescence to account for any artefacts, such as compound interference or aggregation. The resulting competitive binding isotherms were measured and fitted using the expression described by Wang[73] using Pro Fit software package (Quan Soft).

**Isothermal titration calorimetry**

Isothermal titration calorimetry (ITC) was performed using a Microcal itc200 instrument at 25 °C, in the following experimental buffer (unless specifically indicated otherwise): 0.1 M HEPES/Na pH 7.4, 0.1 M magnesium acetate, 0.05 M NaCl, with the addition of 10% v/v DMSO, fresh 1 mM ATP and fresh 0.25 mM TCEP.

Prior the experiment, Aurora A protein was thawed and buffer exchanged in the experimental buffer using NAP-5 Columns (GE Healthcare). Experiments typically involved titrating 25 µM of protein in the sample cell with 300 µM of compound in the syringe. The raw ITC data were fitted using a single site binding model using the Microcal ITC LLC data analysis program in the Origin 7.0 package.

**Crystallisation and structure determination**

To solution of 3.8 mg/ml of Aurora A SilverBullet screen solution 82 (Hampton Research) trans-1,2-cyclohexanedicarboxylic acid was added to final concentration of 8% by volume and the sample was centrifuged for 5 minutes at room temperature at maximum speed in a microcentrifuge. Crystallisation was done in 96-well “MRC” plates (Molecular Dimensions) using a Mosquito nanoliter robot (TTP Labtech) with 300 nl + 300 nl drop with 30% PEG5000 MME (28-32%), 0.1M $(NH_4)_2SO_4$, 0.1 M MES pH 6.5 as the mother liquor. For soaking 1 µl of 100 mM compound in DMSO was diluted with 9 µl of 30% PEG5000 MME (28-32%), 0.1M $(NH_4)_2SO_4$, 0.1 M MES pH 6.5 and added to the crystals for between 2 h and overnight. Crystals were collected into a nylon loop and flash cooled to in liquid nitrogen and stored for data collection. Data collection was typically done for 180 images at 1° oscillation per image. Data reduction and automatic structure determination was done using the pipedream work-flow from Global Phasing Ltd with automatic ligand fitting. Ligand restraints were generated with grade and mogul from CCDC. Structure was analysed and corrected using Coot and refined with Buster TNT. Final ligand electron densities are shown in **Fig. S10**. The structure factors and coordinates have been deposited to Protein Data Bank under access codes: 8C1M, 8C15, 8C1D, 8C1M, 8C15, 8C1D, 8C1H, 8C14, 8C1I, 8C1K with data collection and structrure refinement statistics listed in **Table S4**.

#### Biophysical, cellular and in vivo analyses

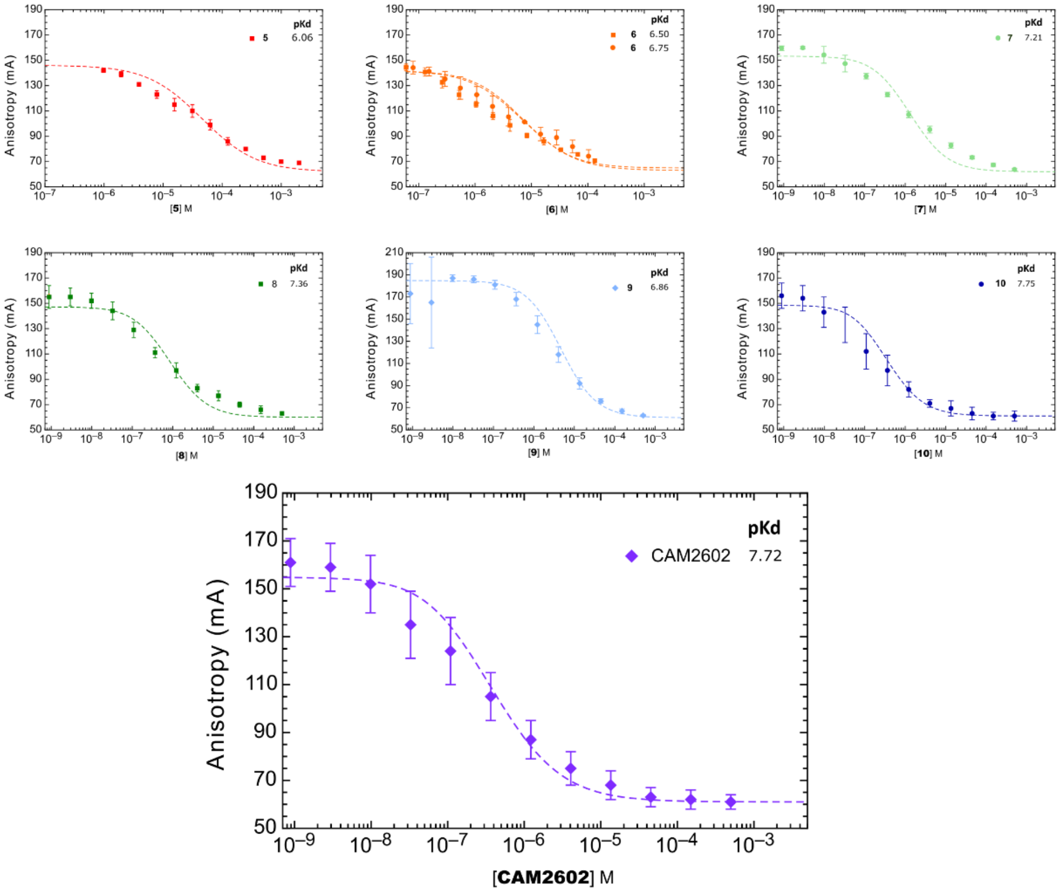

**Figure S1. Fluorescence polarisation anisotropy assay.**
Alexa Fluor 488-labelled TPX2 peptide was displaced from Aurora A by increasing concentration of inhibitors.

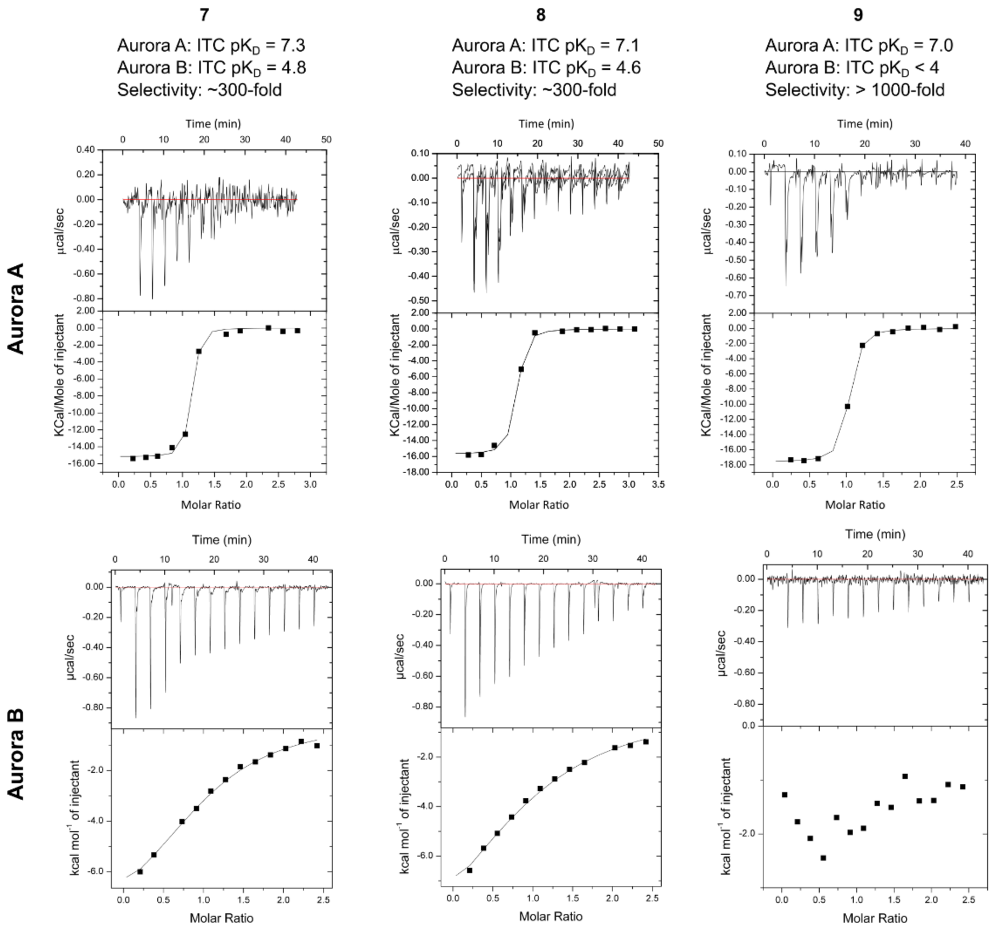

**Figure S2. ITC analysis of selectivity for Aurora A vs. Aurora B for compounds 7,8, and 9.**

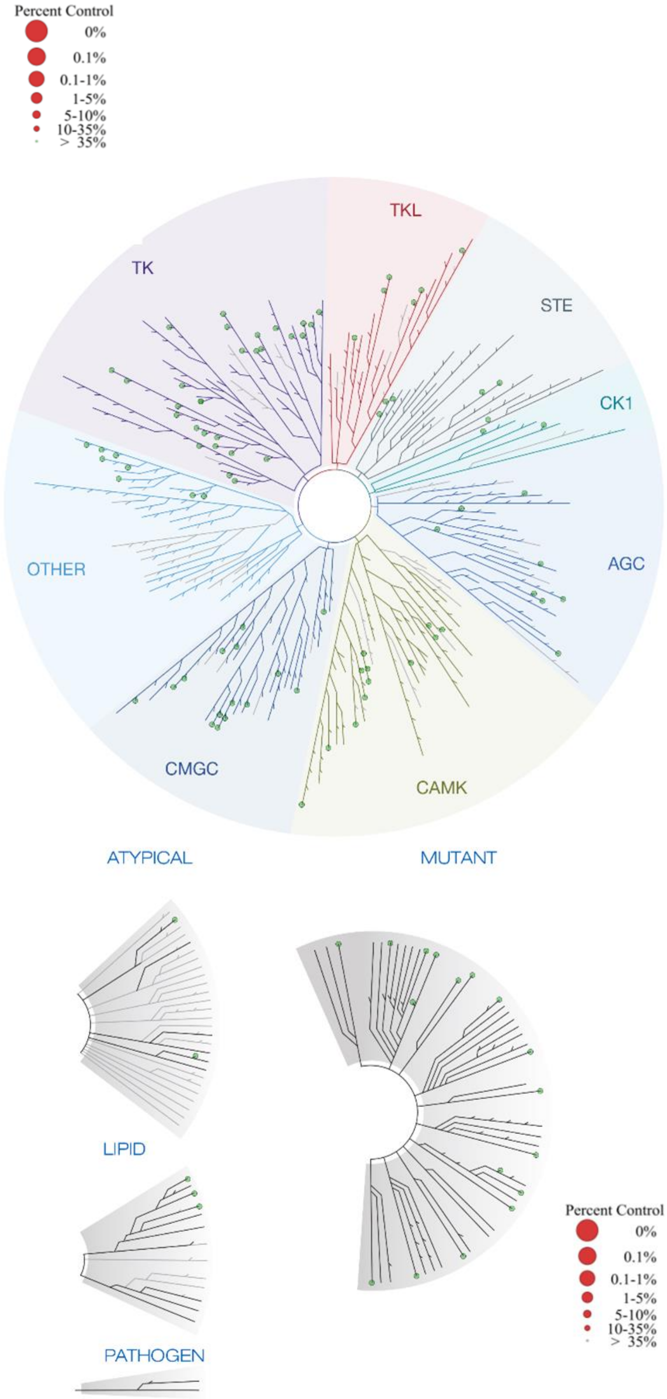

| Target | 8 |
|---|---|
| Gene Symbol | %Ctrl @ 10000nM |
| ABL1(E255K)-phosphorylated | 100 |
| ABL1(T315I)-phosphorylated | 100 |
| ABL1-nonphosphorylated | 99 |
| ABL1-phosphorylated | 100 |
| ACVR1B | 100 |
| ADCK3 | 99 |
| AKT1 | 100 |
| AKT2 | 100 |
| ALK | 100 |
| AURKA | 71 |
| AURKB | 100 |
| AXL | 93 |
| BMPR2 | 100 |
| BRAF | 100 |
| BRAF(V600E) | 100 |
| BTK | 94 |
| CDK11 | 93 |
| CDK2 | 95 |
| CDK3 | 100 |
| CDK7 | 98 |
| CDK9 | 81 |
| CHEK1 | 94 |
| CSF1R | 99 |
| CSNK1D | 100 |
| CSNK1G2 | 91 |
| DCAMKL1 | 96 |
| DYRK1B | 97 |
| EGFR | 99 |
| EGFR(L858R) | 100 |
| EPHA2 | 99 |
| ERBB2 | 100 |
| ERBB4 | 97 |
| ERK1 | 87 |
| FAK | 100 |
| FGFR2 | 94 |
| FGFR3 | 96 |
| FLT3 | 100 |
| GSK3B | 88 |
| IGF1R | 79 |
| IKK-alpha | 100 |
| IKK-beta | 100 |
| INSR | 100 |
| JAK2(JH1domain-catalytic) | 94 |
| JAK3(JH1domain-catalytic) | 98 |
| JNK1 | 93 |
| JNK2 | 83 |
| JNK3 | 80 |
| KIT | 100 |

| Target | 8 |
|---|---|
| Gene Symbol | %Ctrl @ 10000nM |
| KIT(D816V) | 93 |
| KIT(V559D,T670I) | 98 |
| LKB1 | 95 |
| MAP3K4 | 85 |
| MAPKAPK2 | 89 |
| MARK3 | 100 |
| MEK1 | 98 |
| MEK2 | 100 |
| MET | 91 |
| MKNK1 | 98 |
| MKNK2 | 95 |
| MLK1 | 86 |
| p38-alpha | 97 |
| p38-beta | 100 |
| PAK1 | 85 |
| PAK2 | 52 |
| PAK4 | 93 |
| PCTK1 | 100 |
| PDGFRA | 100 |
| PDGFRB | 100 |
| PDPK1 | 96 |
| PIK3C2B | 97 |
| PIK3CA | 100 |
| PIK3CG | 100 |
| PIM1 | 94 |
| PIM2 | 98 |
| PIM3 | 98 |
| PKAC-alpha | 86 |
| PLK1 | 96 |
| PLK3 | 91 |
| PLK4 | 100 |
| PRKCE | 99 |
| RAF1 | 100 |
| RET | 94 |
| RIOK2 | 100 |
| ROCK2 | 100 |
| RSK2(Kin.Dom.1-N-terminal) | 95 |
| SNARK | 95 |
| SRC | 79 |
| SRPK3 | 100 |
| TGFBR1 | 100 |
| TIE2 | 100 |
| TRKA | 85 |
| TSSK1B | 74 |
| TYK2(JH1domain-catalytic) | 100 |
| ULK2 | 100 |
| VEGFR2 | 100 |
| YANK3 | 92 |
| ZAP70 | 100 |

**Figure S3. DiscoverX KINOMEscan specificity screen of compound 9.**
Compound **9** was applied at 10 µM to proprietary cell-free assays, which estimate kinase target inhibition as a product of blocking association with immobilised ATP. (**Left**) TREEspot$^{TM}$ plots of the 97 kinases in the screen organised by kinase family where circles indicate target dendrogram position and size/colour indicate degree of inhibition. (**Right**) List of the 97 kinases screened alongside percent-inhibition detected.

**Table S1. High content toxicology analysis.**

| Cell health parameter | Change | MEC (µM) | $AC_{50}$ (µM) | First signal MEC | First signal $AC_{50}$ |
|---|---|---|---|---|---|
| Cell count | ↓ | 60.7 | >100 | | • |
| Nuclear size | ↑ | 45.5 | >100 | | • |
| DNA structure | ↑ | 29.9 | >100 | | • |
| Cell membrane permeability | ↑ | 10.1 | >100 | • | • |
| Mitochondrial mass | | NR | NR | | |
| Mitochondrial membrane potential | | NR | NR | | |
| Cytochrome c | ↑ | 15.3 | >100 | | • |

Summary of high content toxicology from 72h dose response experiment up to 100 µM of **7** on HepG3 cells. MEC: Minimum effective concentration that significantly crosses vehicle control threshold; $AC_{50}$: The concentration at which 50% maximum effect is observed for each cell health parameter. First Signal: the cell health feature which responds at the lowest observed dose (marked by •). NR: No response observed.

**Table S2. Cerep Express Profile screen data.**

| CEREP panel data | 7 at 10 µM | CAM2602 at 10 µM |
|---|---|---|
| Assay | % Inhibition of Control Specific Binding | % Inhibition of Control Specific Binding |
| A1 (h) (antagonist radioligand) | -11 | 4 |
| A2A (h) (agonist radioligand) | -16 | -17 |
| A3 (h) (agonist radioligand) | 25 | 55 |
| alpha 1 (non-selective) (antagonist radioligand) | 4 | -1 |
| alpha 2 (non-selective) (antagonist radioligand) | -3 | -19 |
| beta 1 (h) (agonist radioligand) | 5 | 6 |
| beta 2 (h) (agonist radioligand) | 6 | 0 |
| AT1 (h) (antagonist radioligand) | 12 | 5 |
| BZD (central) (agonist radioligand) | -2 | -14 |
| B2 (h) (agonist radioligand) | -8 | -73 |
| CB1 (h) (agonist radioligand) | -7 | -26 |
| CCK1 (CCKA) (h) (agonist radioligand) | -24 | -48 |
| D1 (h) (antagonist radioligand) | -1 | 2 |
| D2S (h) (antagonist radioligand) | 2 | 4 |
| ETA (h) (agonist radioligand) | 16 | 21 |
| GABA (non-selective) (agonist radioligand) | -17 | -23 |
| GAL2 (h) (agonist radioligand) | 1 | -1 |
| CXCR2 (IL-8B) (h) (agonist radioligand) | -6 | 1 |
| CCR1 (h) (agonist radioligand) | -8 | -8 |
| H1 (h) (antagonist radioligand) | 4 | -7 |
| H2 (h) (antagonist radioligand) | -6 | -3 |
| MC4 (h) (agonist radioligand) | -1 | -3 |
| MT1 (ML1A) (h) (agonist radioligand) | 0 | 0 |
| M1 (h) (antagonist radioligand) | 1 | 7 |

| | | |
|---|---|---|
| M2 (h) (antagonist radioligand) | -8 | -21 |
| M3 (h) (antagonist radioligand) | 7 | -4 |
| NK2 (h) (agonist radioligand) | 0 | -5 |
| NK3 (h) (antagonist radioligand) | -3 | -20 |
| Y1 (h) (agonist radioligand) | 4 | -1 |
| Y2 (h) (agonist radioligand) | -1 | -6 |
| NTS1 (NT1) (h) (agonist radioligand) | -19 | -1 |
| delta (DOP) (h) (agonist radioligand) | 2 | 14 |
| kappa (KOP) (agonist radioligand) | 7 | 7 |
| mu (MOP) (h) (agonist radioligand) | 1 | 5 |
| NOP (ORL1) (h) (agonist radioligand) | 17 | 0 |
| EP4 (h) (agonist radioligand) | 14 | 4 |
| 5-HT1A (h) (agonist radioligand) | -4 | 20 |
| 5-HT1B (antagonist radioligand) | 1 | -1 |
| 5-HT2A (h) (antagonist radioligand) | 0 | -11 |
| 5-HT2B (h) (agonist radioligand) | 18 | 22 |
| 5-HT3 (h) (antagonist radioligand) | 9 | 7 |
| 5-HT5a (h) (agonist radioligand) | 12 | 0 |
| 5-HT6 (h) (agonist radioligand) | 3 | 0 |
| 5-HT7 (h) (agonist radioligand) | 11 | 3 |
| sst (non-selective) (agonist radioligand) | 2 | 1 |
| VPAC1 (VIP1) (h) (agonist radioligand) | -7 | -7 |
| V1a (h) (agonist radioligand) | 4 | 3 |
| Ca2+ channel (L, verapamil site) (phenylalkylamine) (antagonist radioligand) | -12 | -5 |
| KV channel (antagonist radioligand) | 0 | -3 |
| SKCa channel (antagonist radioligand) | 3 | -2 |
| Na+ channel (site 2) (antagonist radioligand) | -10 | -5 |
| Cl- channel (GABA-gated) (antagonist radioligand) | 13 | 28 |
| norepinephrine transporter (h) (antagonist radioligand) | -2 | 6 |
| dopamine transporter (h) (antagonist radioligand) | 13 | 11 |
| 5-HT transporter (h) (antagonist radioligand) | -9 | -5 |

**7** and **CAM2602** were screened at 10 µM against a panel of 55 GPCRs, transporters and ion channels. The percentage inhibition of the binding of a radioactively labelled ligand specific for each target is given.

**Table S3. Calculated and measured ADMET properties for CAM2602**

| Property | Value |
| ---: | --- |
| MW | 483.97 Da |
| TPSA | 96.9 Å$^2$ |
| LogP[a] | 4.38 |
| $pK_D$ (FP) | 7.4 |
| Microsomal CLint | Mouse $t_{1/2}$ = 106 min |
| | 13.1 µL/min/mg protein |
| | Human $t_{1/2}$ = 155 min |
| | 8.9 µL/min/mg protein |
| Hepatocyte CLint | Mouse $t_{1/2}$ = 139 min |
| | 10 µL/min/$10^{-6}$ cells |
| | Rat $t_{1/2}$ = 408 min |
| | 3.4 µL/min/$10^{-6}$ cells |
| | Human $t_{1/2}$ = 205 min |
| | 6.8 µL/min/$10^{-6}$ cells |
| hERG inhibition $IC_{50}$[b] | >25 µM |
| Mouse plasma protein binding | 98.5% |
| Human plasma protein binding | 99.6% |
| Caco-2 permeability: | |
| Papp ($10^{-6}$ cm $s^{-1}$) A2B / B2A / Efflux ratio | 5.1 / 41 / 8.1 |
| CYP450 $IC_{50}$[c] | All >25 µM |

[a] Calculated with ChemDraw version 19.1.0.8

[b] Whole-cell voltage clamping assay with mammalian cells transfected with hERG potassium channel

[c] CYP1A (ethoxyresorufin), CYP2C9 (tolbutamide), CYP219 (mephenytoin), CYP2D6 (dextromethorphan), CYP3A4 (midazolam), CYP3A4 (testosterone).

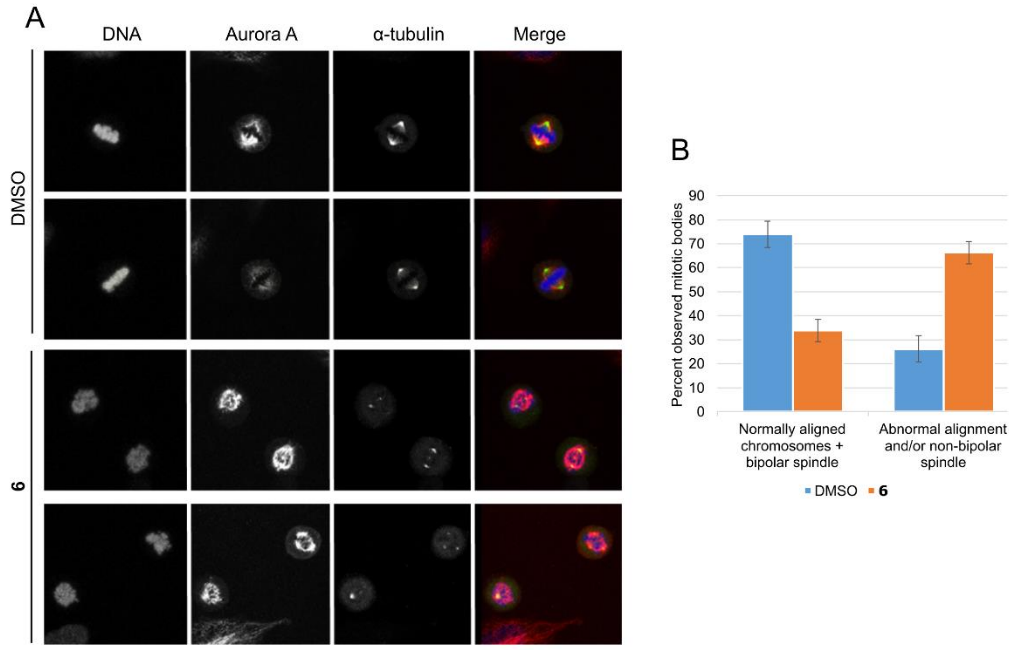

**Figure S4. Mitotic spindle abnormalities in cells treated with 6.**

(**A**) HeLa cells were treated with 50 µM (1x $GI_{50}$) **6** or DMSO for 6 hours prior to being fixed, fluorescently stained for the indicated proteins or DNA and imaged using confocal microscopy. Two representative fields containing mitotic cells are shown for both treatment conditions. Mitotic cells were enumerated to exhibit spindle abnormalities if they demonstrated unaligned chromosomes and/or non-bipolarity, examples of which are indicated by the solid and outline arrowhead, respectively. (**B**) Relative proportions of normal and abnormal spindle classes across all imaged mitotic cells for both DMSO and **5** treated cells (>100 mitotic cell observations). Error bars show standard deviations from the mean (n=3 image sets per condition).

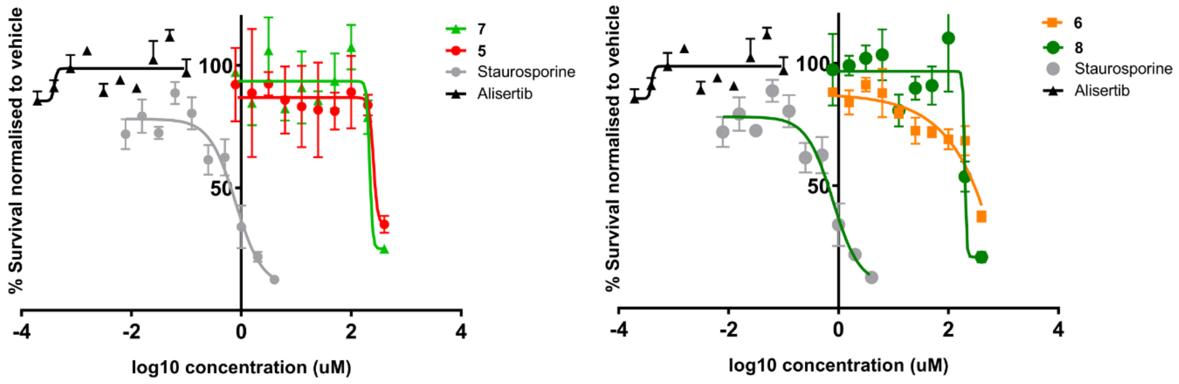

**Figure S5. Toxicity in non-cycling cells.**

Viable, non-cycling peripheral blood mononuclear cells were grown for 72 hours in the presence of increasing concentrations of the indicated compounds, Alisertib or staurosporine as a positive control. After 72 hours the cell media was supplemented with CellTiter Blue dye to fluorescently quantify the viable cells under each treatment condition. Viability values were normalised against concurrently performed vehicle controls (DMSO for CAM and Alisertib, EtOAc for staurosporine). Concentrations of compounds **5-8;** 0.8-400 uM; Alisertib 0.2-100 nM; staurosporine 8 nM- 4 uM.

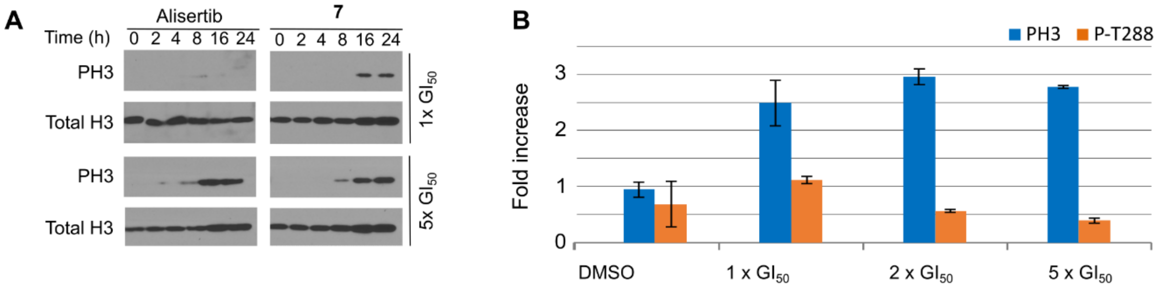

**Figure S6. PH3 levels evaluation with compound 7.**
**(A)** Western blot analysis of PH3 levels in Jurkat cells treated with the indicated fold-$GI_{50}$ equivalents of alisertib (7 or 35 nM) and **7** (20 or 100 µM). **(B)** Flow cytometric analysis of Jurkat cells treated with a range of fold-$GI_{50}$ concentrations of **7** (1x $GI_{50}$ = 20 µM) for 8 h. The cells were stained for DNA, PH3 and P-Thr288 Aurora A and were analysed to determine the proportion of mitotic cells (having both 4n DNA and PH3 positivity); additionally, the proportion of cells positive for P-Thr288 within the mitotic population was also measured per treatment condition. Data is plotted as normalised values relative to the untreated control.

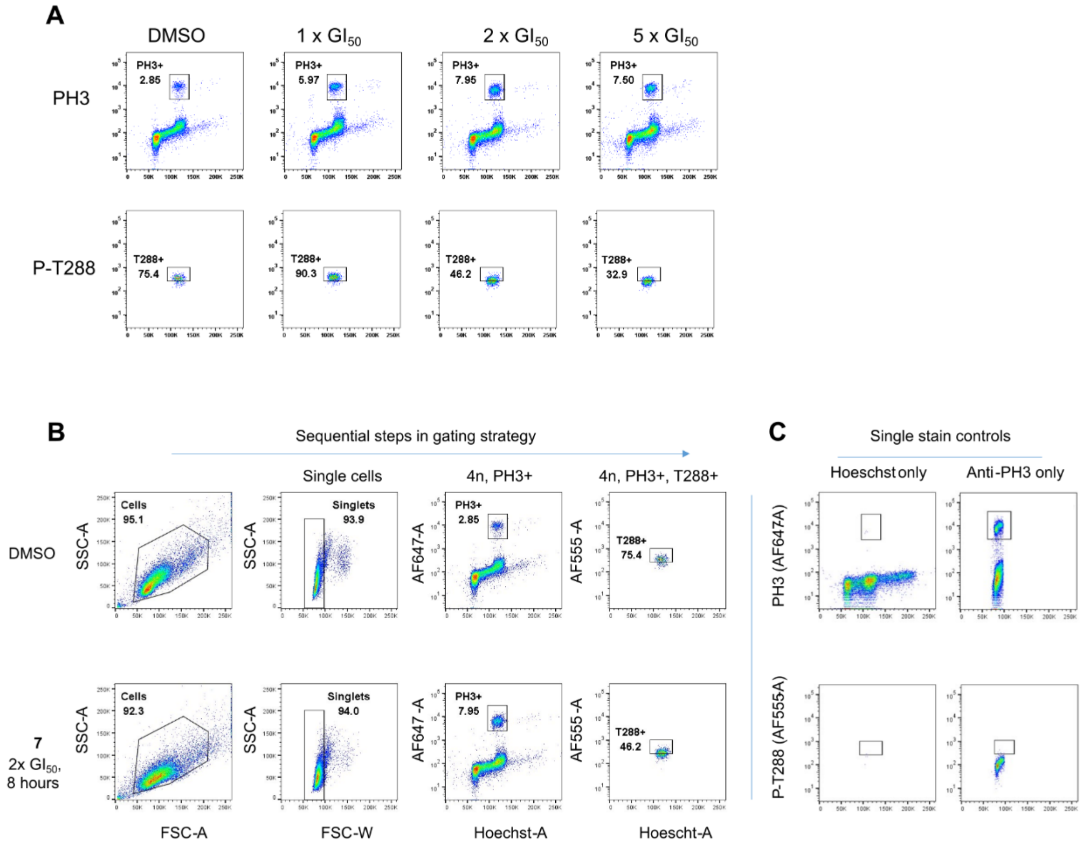

**Figure S7. Flow cytometry gating strategy to detect Aurora A inhibition biomarkers.**
(A) Flow cytometric data as summarised in main **Fig. 5**. (B) Flow cytometric analysis of Jurkat cells treated for 8 hours with either DMSO or 2x $GI_{50}$ concentrations of **7**. From left to right, the four scatter plots show the sequential analysis used to detect the Aurora A biomarker changes resulting from the compound treatment applied using flow cytometry. Each scatter plot represents only those cells within the gated population (box) of the previous plot to the left. The leftmost two panels

concern the gating of non-debris, single cells in the samples through light scattering properties. The third panel plots these singlets against Hoechst-staining intensity of DNA (X-axis) and Alexa Fluor-labelled anti-phospho-histone 3 (PH3) antibody intensity (Y-axis). The gated population is both PH3-positive and possess 4n DNA, as expected from mitotic cells. In the final panel, the PH3+,4n cells are gated if positive for T288-phosphorylated (P-T288) Aurora A. (C) Positions of the gates for PH3 and P-T288 positivity were determined in untreated, single stain samples of cells.

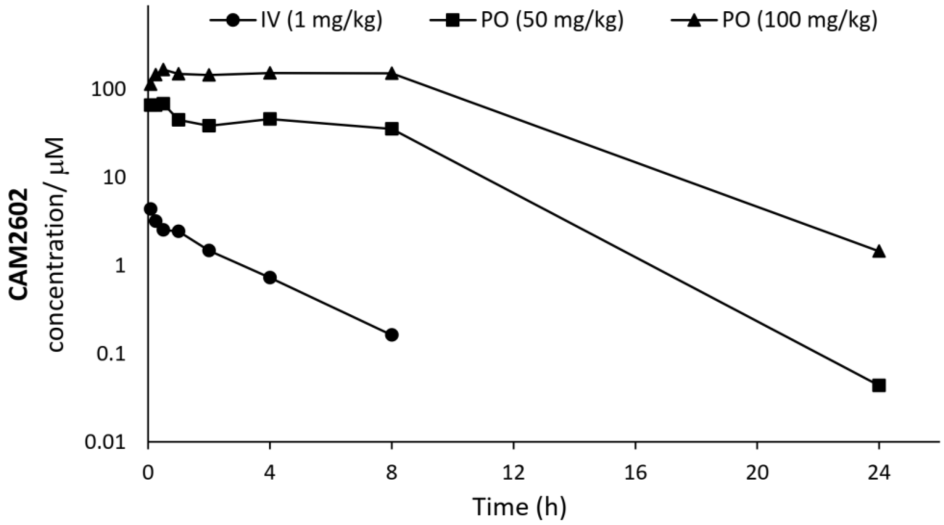

**Figure S8. Pharmacokinetics of CAM2602.**
**CAM2602** was administered at three separate doses in female CD-1 mice and measuring the total concentration of compound in plasma over time.

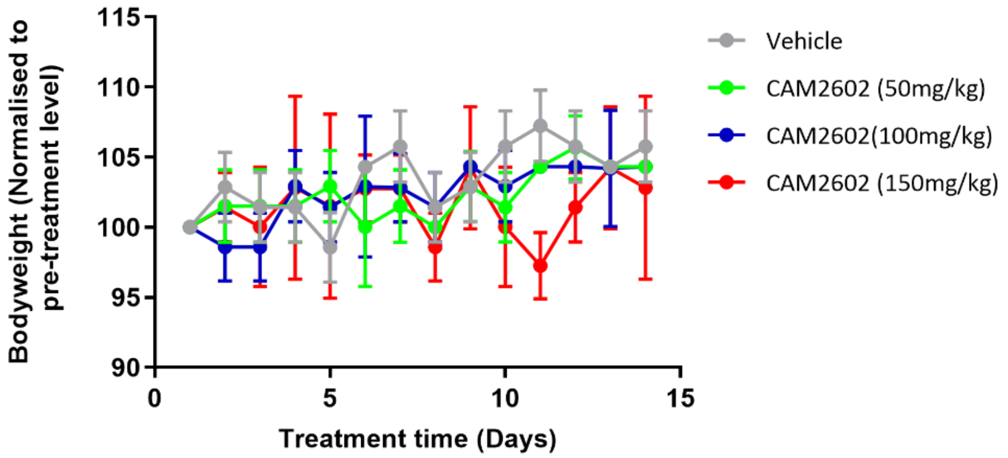

**Figure S9. Tolerability of CAM2602.**
The indicated doses of CAM2602 were orally administered to female NSG mice, QD for 7 days, followed by 7 days with no dosing. Plotted are normalised bodyweight measurements taken daily. Values shown are mean ±SD; n=3 for all groups. Compound formulation: DMSO:20% HP-β-CD (2-hydroxypropyl-beta-cyclodextrin) in PBS (2.5:97.5) pH 7.6. At all doses, no ill-health indications were observed.

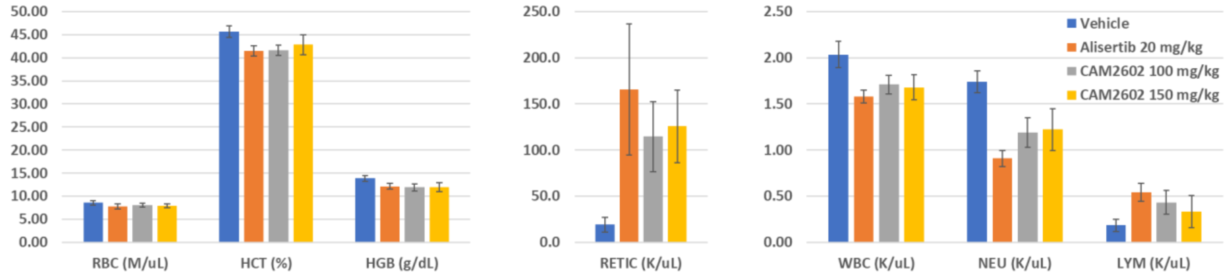

**Figure S10. Analysis of blood samples upon conclusion of efficacy study.**

On the final day of the study, blood samples from each mouse were assessed to determine relative population sizes of each blood cell type per sample. Units are shown adjacent to categories: absolute counts (millions or thousands per microliter, M/µl, K/µl) or weight per volume (g/dL) or percentage parent population, where indicated. Categories: red blood cells (RBC); haematocrit (HCT); haemoglobin (HGB); reticulocytes (RETIC); white blood cells (WBC); neutrophils (NEU); lymphocytes (LYM). Values are means and standard deviations taken from 5 biological replicates.

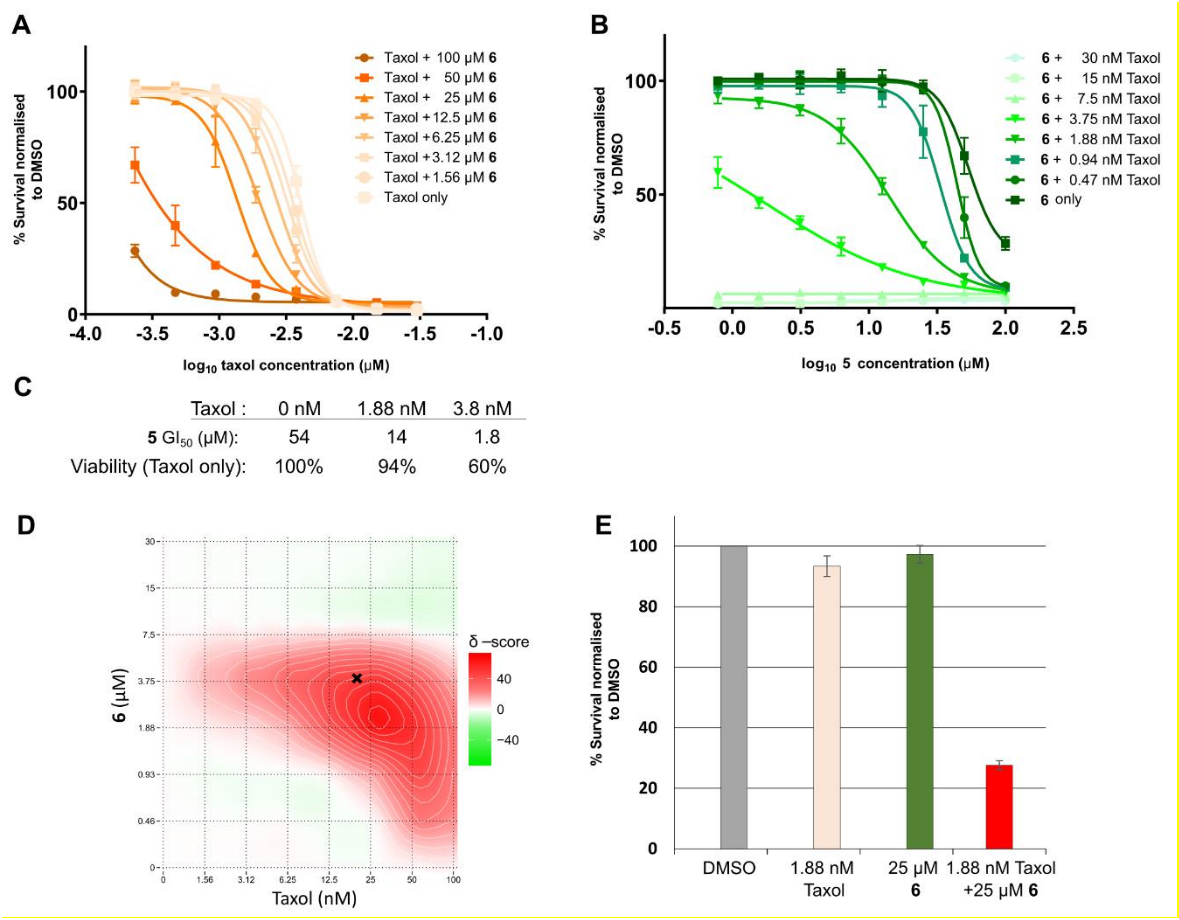

| Taxol : | 0 nM | 1.88 nM | 3.8 nM |
|---|---|---|---|
| **5** $GI_{50}$ (μM): | 54 | 14 | 1.8 |
| Viability (Taxol only): | 100% | 94% | 60% |

Figure S11. Aurora A:TPX2 PPI inhibitors synergise with Taxol in PANC-1 cells.
**(A)** and **(B)**: PANC-1 cells were dosed with a matrix of concentrations of Taxol and **6**, including single agent and vehicle controls for all concentrations tested. 72 hours following treatment, the cells were assayed for remaining viability relative to vehicle controls. **(C)** Table showing effective decrease in **6** $GI_{50}$ in PANC-1 cells when combined with increasing concentrations of Taxol. Also shown are the corresponding viability changes effected by Taxol if applied as a single agent. (**D**) The vehicle-normalised viability assay data were processed using SynergyFinder webserver (https://synergyfinder.org/)[57], producing a heatmap indicating the presence of synergy (red) or antagonism (green) between the two drugging agents when compared to modelled predictions of additivity (**E**) Chart comparing vehicle-normalised 72-hour viability assay values between single agent and combined treatments of the concentrations of Taxol and **6** yielding the greatest synergic effect. The single-agent inhibition values for Taxol alone or **6** alone were used to calculate a drug combination surface under the assumption of an additive effect using SynergyFinder, which is shown as the 'predicted' value. Bars show standard deviations from the mean (n=4).

**Table S4. Crystallographic data collection and refinement parameters.**

| Ligand | 2 | 3 | 4 |
|---|---|---|---|
| PDB code | 8C1M | 8C15 | 8C1D |
| **Data Collection:** | | | |
| Beamline | DLS I04-1 | ESRF ID29 | ESRF MASSIF-3 |
| Wavelength (Å) | 0.92 | 0.973 | 0.9677 |
| Resolution range (Å) | 2.84-53.97 (2.91-2.84) | 70.02 - 2.41 (2.416 - 2.407) | 70.44 - 2.12 (2.122 - 2.115) |
| Space group | P 41 21 2 | P 61 2 2 | P 61 2 2 |
| Cell (a b c) (Å) | 82.82 82.82 139.0 | 80.86 80.86 165.16 | 81.34 81.34 171.53 |
| Cell (α β γ) (°) | 90.00 90.00 90.00 | 90.00 90.00 120.00 | 90.00 90.00 120.00 |
| Total reflections | 148180 (11280) | 227396 (2326) | 435735 (4149) |
| Unique reflections | 11913 (847) | 13083 (129) | 19797 (178) |
| Multiplicity | 12.4 (13.3) | 17.4 (18.0) | 22.0 (23.3) |
| Completeness (%) | 100.0 (99.8) | 100.0 (98.5) | 99.2 (96.2) |
| Mean I/σ(I) | 25.5 (3.7) | 16.6 (1.6) | 23.5 (1.3) |
| $R_{merge}$ | 0.071 (0.828) | 0.103 (1.94) | 0.069 (2.90) |
| $R_{pim}$ | 0.028 (0.323) | 0.026 (0.46) | 0.015 (0.61) |
| CC ½ | 1.000 (0.963) | 0.999 (0.67) | 0.999 (0.67) |
| **Refinement:** | | | |
| R / $R_{free}$ | 0.2529/0.2620 | 0.199 / 0.207 | 0.210 / 0.238 |
| No. of atoms | 2150 | 2251 | 2291 |
| No of ligand atoms | 50 | 102 | 103 |
| No of waters | 0 | 40 | 83 |
| Number of protein residues | 262 | 259 | 259 |
| Average/Wilson B factor (Å$^2$) | 82.353/72.37 | 75.6 / 74.1 | 67.0 / 61.4 |
| B-factor for ligands (Å$^2$) | 100.2 | 99.4 | 92.1 |
| B-factor for solvent (Å$^2$) | n/a | 71.1 | 68.9 |
| RMS (bonds) (Å) | 0.006 | 0.01 | 0.009 |
| RMS (bond angles) (°) | 0.87 | 1.08 | 1.03 |
| RMS (dihedral angles) (°) | 2.55 | 3.02 | 3.04 |

| Ligand | 5 | 6 | 7 |
|---|---|---|---|
| PDB code | 8C1E | 8C1F | 8C1G |
| **Data Collection:** | | | |
| Beamline | SOLEIL PROXIMA 2 | ESRF MASSIF-3 | DLS I03 |
| Wavelength (Å) | 0.9801 | 0.9677 | 0.97 |
| Resolution range (Å) | 135.87 - 2.80 (2.808 - 2.798) | 65.26 - 1.92 (1.931 - 1.924) | 53.33 - 1.96 (1.965 - 1.959) |
| Space group | P 41 21 2 | P 61 2 2 | P 61 2 2 |
| Cell (a b c) (Å) | 82.04 82.04 135.87 | 81.44 81.44 172.01 | 80.82 80.82 164.67 |
| Cell (α β γ) (°) | 90.00 90.00 90.00 | 90.00 90.00 120.00 | 90.00 90.00 120.00 |
| Total reflections | 83838 (974) | 405731 (3313) | 224440 (2256) |
| Unique reflections | 12082 (135) | 24878 (216) | 22054 (234) |
| Multiplicity | 6.9 (7.2) | 16.3 (15.3) | 10.2 (9.6) |
| Completeness (%) | 100.0 (100.0) | 94.2 (83.7) | 93.2 (98.7) |
| Mean I/σ(I) | 14.0 (3.4) | 25.8 (1.7) | 15.3 (1.7) |
| $R_{merge}$ | 0.167 (1.16) | 0.057 (1.56) | 0.075 (0.87) |
| $R_{pim}$ | 0.069 (0.47) | 0.014 (0.41) | 0.025 (0.29) |
| CC ½ | 0.994 (0.55) | 1.000 (0.69) | 0.999 (0.81) |
| **Refinement:** | | | |
| R / $R_{free}$ | 0.173 / 0.239 | 0.205 / 0.234 | 0.199 / 0.225 |
| No. of atoms | 2279 | 2329 | 2326 |
| No of ligand atoms | 54 | 131 | 120 |
| No of waters | 73 | 90 | 115 |
| Number of protein residues | 262 | 258 | 258 |
| Average/Wilson B factor (Å$^2$) | 48.8 / - | 49.1 / 42.1 | 45.2 / 36.4 |
| B-factor for ligands (Å$^2$) | 41.3 | 71.5 | 59.5 |
| B-factor for solvent (Å$^2$) | 47.2 | 50.5 | 48.2 |
| RMS (bonds) (Å) | 0.015 | 0.01 | 0.01 |
| RMS (bond angles) (°) | 2.102 | 1.09 | 1.06 |
| RMS (dihedral angles) (°) | 7.153 | 3.53 | 3.24 |

| Ligand | 8 | 9 | 10 |
|---|---|---|---|
| PDB code | 8C1H | 8C14 | 8C1I |
| **Data Collection:** | | | |
| Beamline | DLS I03 | ESRF MASSIF-3 | ESRF ID29 |
| Wavelength (Å) | 0.97 | 0.9677 | 0.973 |
| Resolution range (Å) | 64.61 - 2.23 (2.241 - 2.233) | 81.04 - 1.93 (1.937 - 1.930) | 83.63 - 2.81 (2.820 - 2.810) |
| Space group | P 61 2 2 | P 41 21 2 | P 61 2 2 |
| Cell (a b c) (Å) | 81.03 81.03 165.62 | 81.04 81.04 138.18 | 81.43 81.43 167.26 |
| Cell (α β γ) (°) | 90.00 90.00 120.00 | 90.00 90.00 90.00 | 90.00 90.00 120.00 |
| Total reflections | 184733 (1826) | 418805 (4053) | 149762 (1686) |
| Unique reflections | 15901 (152) | 32771 (332) | 8604 (93) |
| Multiplicity | 11.6 (12.0) | 12.8 (12.2) | 17.4 (18.1) |
| Completeness (%) | 97.4 (98.7) | 92.7 (100.0) | 100.0 (98.9) |
| Mean I/σ(I) | 13.6 (1.9) | 15.2 (1.4) | 17.0 (1.4) |
| $R_{merge}$ | 0.089 (0.98) | 0.109 (2.23) | 0.148 (1.91) |
| $R_{pim}$ | 0.027 (0.28) | 0.032 (0.66) | 0.036 (0.45) |
| CC ½ | 0.999 (0.86) | 0.999 (0.60) | 0.998 (0.72) |
| **Refinement:** | | | |
| R / $R_{free}$ | 0.211 / 0.245 | 0.181 / 0.194 | 0.212 / 0.234 |
| No. of atoms | 2294 | 2513 | 2249 |
| No of ligand atoms | 113 | 128 | 111 |
| No of waters | 77 | 205 | 14 |
| Number of protein residues | 258 | 264 | 260 |
| Average/Wilson B factor (Å$^2$) | 63.4 / 53.5 | 42.9 / 38.2 | 81.5 / 91.9 |
| B-factor for ligands (Å$^2$) | 77.3 | 67.7 | 103.7 |
| B-factor for solvent (Å$^2$) | 63.4 | 54.6 | 65.3 |
| RMS (bonds) (Å) | 0.011 | 0.01 | 0.008 |
| RMS (bond angles) (°) | 1.09 | 1.01 | 1.02 |
| RMS (dihedral angles) (°) | 3.28 | 3.34 | 3.01 |

| Ligand | **CAM2602** |
|---|---|
| PDB code | 8C1K |
| **Data Collection:** | |
| Beamline | DLS I24 |
| Wavelength (Å) | 0.9686 |
| Resolution range (Å) | 169.18 - 2.27 (2.304 - 2.265) |
| Space group | P 61 2 2 |
| Cell (a b c) (Å) | 82.14 82.14 169.18 |
| Cell (α β γ) (°) | 90.00 90.00 120.00 |
| Total reflections | 308579 (4402) |
| Unique reflections | 12671 (190) |
| Multiplicity | 24.4 (23.2) |
| Completeness (%) | 92.5 (65.5) |
| Mean I/σ(I) | 28.3 (1.2) |
| $R_{merge}$ | 0.074 (2.85) |
| $R_{pim}$ | 0.015 (0.60) |
| CC ½ | 1.000 (0.66) |
| **Refinement:** | |
| R / $R_{free}$ | 0.217 / 0.242 |
| No. of atoms | 2264 |
| No of ligand atoms | 115 |
| No of waters | 39 |
| Number of protein residues | 262 |
| Average/Wilson B factor (Å$^2$) | 82.2 / 82.1 |
| B-factor for ligands (Å$^2$) | 104.5 |
| B-factor for solvent (Å$^2$) | 74.9 |
| RMS (bonds) (Å) | 0.011 |
| RMS (bond angles) (°) | 1.25 |
| RMS (dihedral angles) (°) | 3.51 |

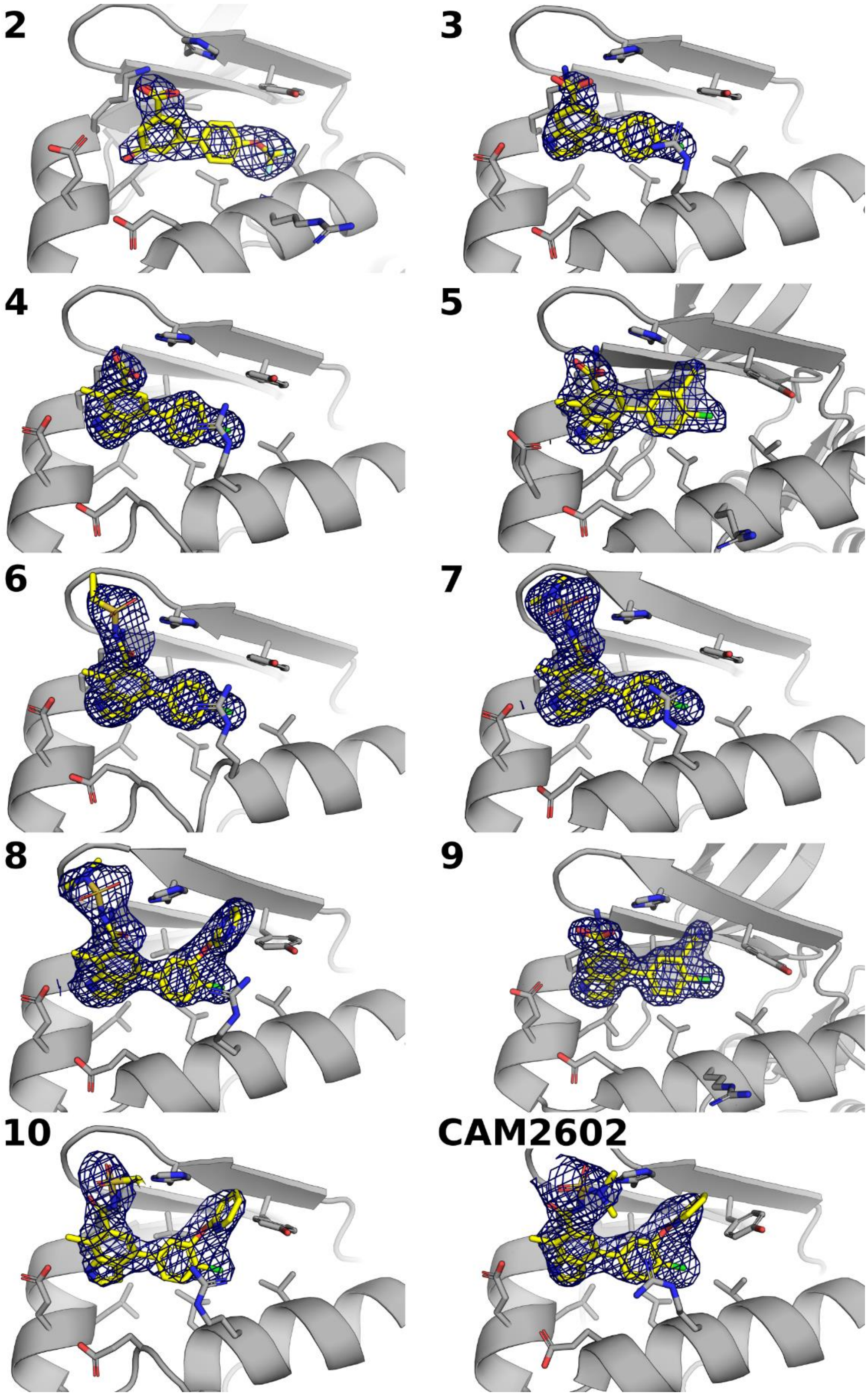

**Figure S12. Electron densities of ligands**

Electron densities of ligands **2-10** and **CAM2602** in complex with Aurora A after final refinement, all contoured at 1σ level.

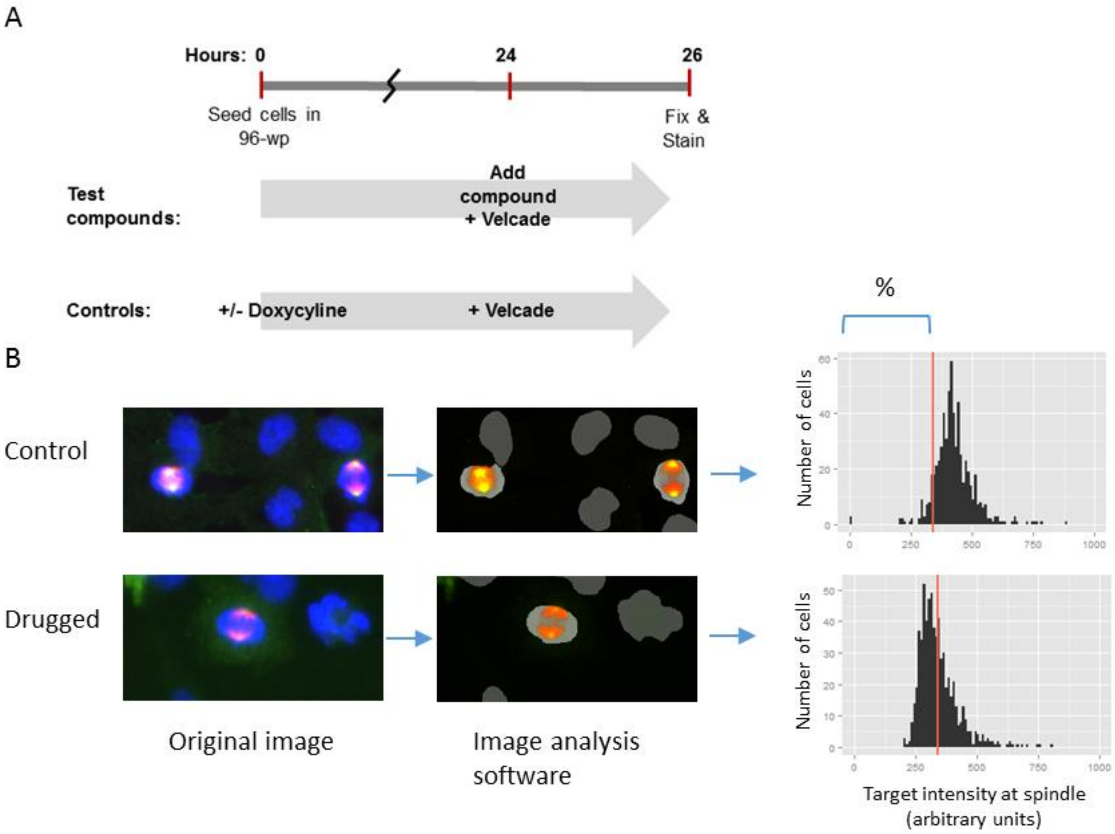

**Figure S13. High-content assay to assess cellular Aurora A engagement by PPI inhibitors.**
(A) Experimental structure for the HCS assay – timelines proceed from seeding cells to 96-well plates through to the final 2 hour treatment of the cells with compound titrations in the presence of Velcade. The control wells of each plate received either DMSO or doxycycline (for mCherry-TPX2-1–43 peptide induction) immediately following cell seeding, 24 hours prior to the drugging of the compound wells. All wells receive Velcade in the last 2 hours of the assay. (B) Example Aurora A mislocalisation assay images of vehicle control and compound treated HeLa cells after 2 hours of treatment. Left panels show merged-channel images of the fixed cells fluorescently stained for DNA (blue), TPX2 (red) and Aurora A (green). The middle panels show the Aurora A and TPX2 fluorescent stains superimposed onto the software-generated mask of nuclei positions generated from the DNA fluorescence data. Only the TPX2-positive, mitotic nuclei are retained for phenotypic assessment. The right panels show histograms of the two example cell populations where the x-axis shows the observed fluorescence intensity values for Aurora A immunostaining within the limits of the TPX2-immunostained mitotic spindle for every mitotic cell. The red line represents the assay threshold, calculated per plate, to the left of which cells exhibit loss of Aurora A from the spindle.

#### Chemical structures of key compounds

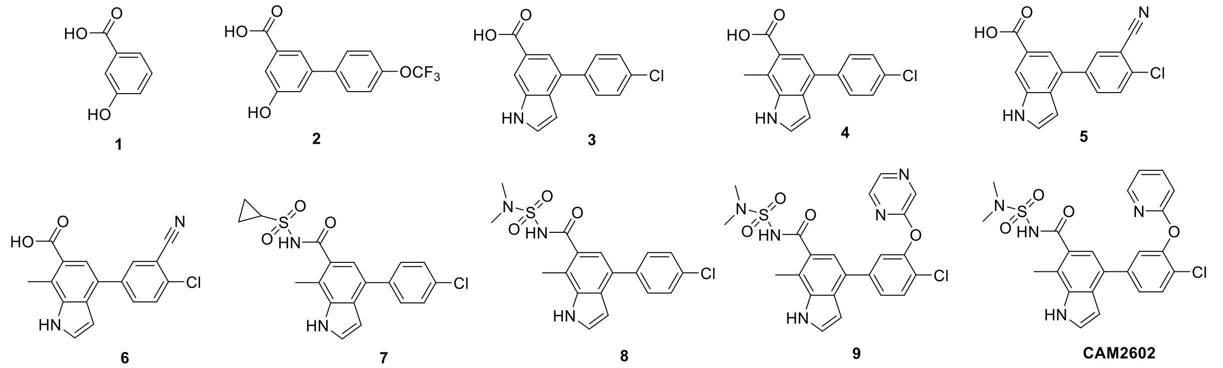

**Supplementary Figure S11** Chemical structures of key compounds

#### Synthetic chemistry schemes

General chemistry information

**Abbreviations:** TEA: triethylamine, DCM: dichloromethane, DME: dimethoxyethane, CDI: carbonyldiimidazole, DBU: 1,8-Diazabicyclo[5.4.0]undec-7-ene, LCMS: liquid chromatography-mass spectrometry

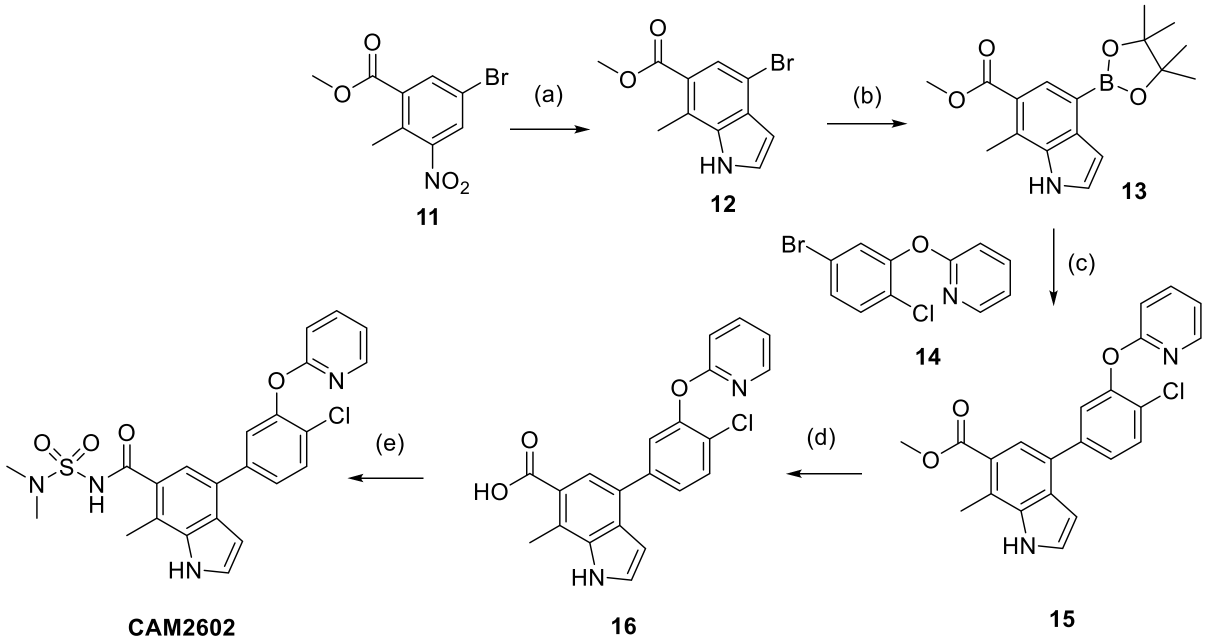

**Scheme S1 Synthesis of compound CAM2602**
(a) vinylmagnesium bromide, THF, -78 °C (b) bis(pinacolato)diboron, Pd(dppf)$Cl_2$·DCM, DMSO, 90 °C, 4 h (c) **14**, Pd(dppf)$Cl_2$ DCM, DME, $H_2O$, 120 °C (microwave), 0.5 h (d) LiOH, THF, water, 45 °C (e) (i) CDI, DBU, THF, 45 °C, 3h (ii) dimethylsulfamide, DBU, 80 °C.

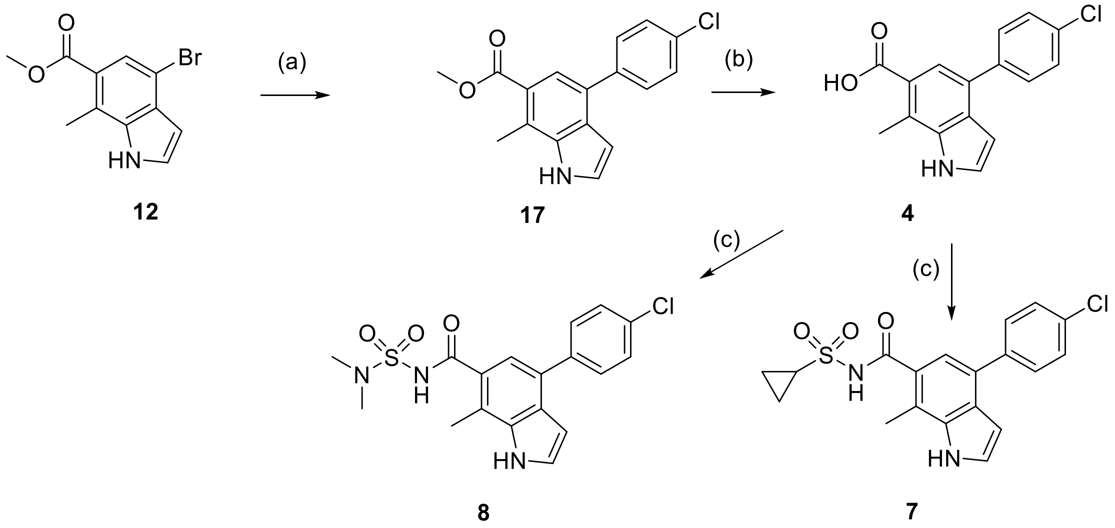

**Scheme S2 Synthesis of compound 6, 7 and 4**

(a) 4-Chlorophenylboronic acid, TEA, DME, water, Pd(dppf)$Cl_2$.$CH_2Cl_2$, 120 °C (microwave), 0.5 h (b) Lithium iodide, 180 °C (microwave), 1 h (c) (i) CDI, DBU, THF, 45 °C, 3h (ii) sulfonamide, DBU, 80 °C.

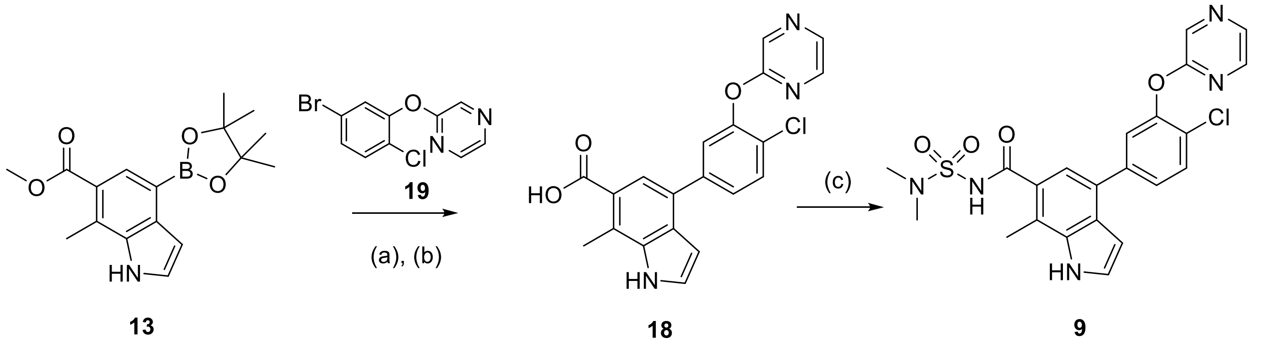

**Scheme S3 Synthesis of compound 18 and 8**

(a) **19**, TEA, DME, water, Pd(dppf)$Cl_2$·$CH_2Cl_2$, 120 °C (microwave), 0.5 h (b) LiOH, THF, water, 45 °C (c) (i) CDI, DBU, THF, 45 °C, 3h (ii) dimethylsulfamide, DBU, 80 °C.

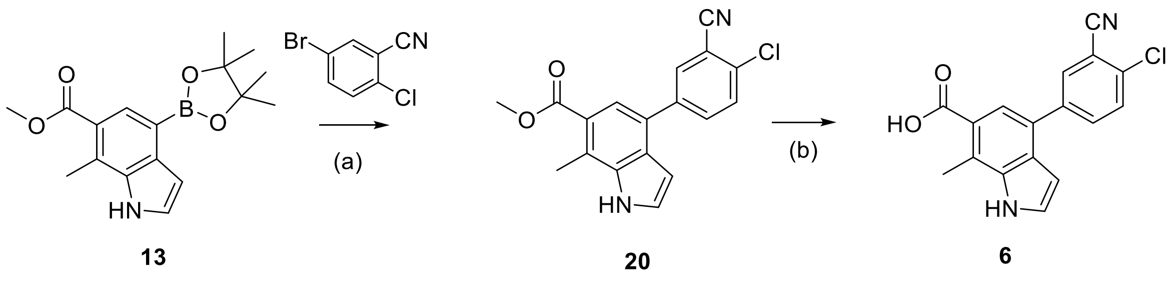

**Scheme S4 Synthesis of compound 5**

(a) 4-chloro-3-cyanophenylboronic acid, TEA, DME, water, Pd(dppf)$Cl_2$·$CH_2Cl_2$, 120 °C, 0.5 h (b) Lithium iodide, 180 °C (microwave), 1 h.

#### Synthetic chemistry methods

##### Method A – Suzuki coupling

Arylbromide (1 equiv.), boronic acid (1 equiv.) and triethylamine (3 equiv.) were dissolved in DME (1.5 ml) and water (0.5 ml) and nitrogen was bubbled through for 10 minutes. Pd(dppf)$Cl_2$·$CH_2Cl_2$ (10 mol%) was added and the reaction was heated with microwave irradiation at 120 ˚C for 30 min. After cooling to room temperature, the solvents were evaporated *in vacuo*. The crude residue was dissolved in DCM (10 ml) filtered through a hydrophobic frit and then evaporated and purified by FC ($SiO_2$, 10-100% EA in pet ether 40-60) to give the product.

##### Method B –Ester hydrolysis, thermal

The methyl ester (1 equiv.) was dissolved in THF (2 mL) and $H_2O$ (2 mL) and LiOH (3 equiv.) was added and stirred overnight at 45 $^{o}$C. After cooling to room temperature, ethyl acetate (2 x 50 mL) and $H_2O$ (50 mL) was added and the organic layers discarded. The aqueous layer was acidified with dilute HCl to pH 4 and extracted with ethyl acetate (2 x 50 mL), dried with $Na_2SO_4$ and the solvent removed *in vacuo* to give the product.

##### Method C – Ester hydrolysis, microwave

The methyl ester (1 equiv.) and lithium iodide (10 equiv.) were dissolved in pyridine (2 mL) and the reaction mixture was heated at 180 ˚C for 1 hour under microwave irradiation. After cooling to room temperature, the solvent was removed *in vacuo* and the residue taken up in ethyl acetate (20 mL) and sat. aq. $NaHCO_3$ (20 mL). The aqueous layer was carefully acidified (pH 2) and extracted with ethyl acetate (2 x 30 mL). The organic layers were combined and the solvent removed *in vacuo* to give the product.

##### Method D1 – Biaryl ether formation

To a stirred solution of phenol (1.5 mmol) in DMF (1.3mL) was added potassium carbonate (2.5 mmol) and the appropriate 2-bromopyridine (1.5 mmol). The reaction mixture was stirred at 150 °C overnight. The mixture was diluted in water and the organic layer was extracted with EtOAc (x3). The combined organic layers were washed with brine and dried over $Na_2SO_4$ and filtered. The filtrate was evaporated *in vacuo* to obtain the crude which was purified by FC ($SiO_2$, 0-25 % EA in pet ether 40-60) to provide the product.

##### Method D2 – Biaryl ether formation

A stirred solution of phenol (1-1.5 mmol) and Caesium Carbonate (2.5 mmol) in dry DMSO (5ml) was heated at 45°C for 10 minutes. The appropriate fluoro-pyrimidine (1 mmol) or fluoro-pyrazine was then added to the mixture, the mixture was flushed with nitrogen and heated between 65°C and 150°C in sealed microwave vials for 1-16h according to the starting materials reactivity. The resulting mixture was poured into water and extracted with EtOAc (3 x). The combined organics were dried over $Na_2SO_4$ and filtered. The filtrate was evaporated *in vacuo* to give a crude product which was purified by FC ($SiO_2$, 0-25 % EtOAc in pet ether 40-60) to provide the product.

##### Method E – Sulfonamide coupling

A solution of the requisite carboxylic acid (0.21 mmol) and carbonyldiimidazole (0.63 mmol, 3.0 eq) in THF (6 mL) was heated at 45°C for 3 h. Then a solution of DBU (0.84 mmol, 4.0 eq) and the requisite sulfonamide (0.31 mmol, 1.5 eq) in THF (2 mL) was added and stirring continued at 80°C overnight. If the reaction was incomplete after 18 h, additional sulfonamide was added and stirred overnight. Upon completion the reaction mixture was concentrated *in vacuo*, diluted with DCM:IPA (4:1, 50 mL), washed with 1M HCl (2 × 25 mL) and brine (25 mL) then passed through a hydrophobic frit and solvent removed *in vacuo*. Purification via FC ($SiO_2$, 10-60% EtOAc in pet ether 40-60 (both with 0.5% AcOH)) provided the desired product.

#### Chemical synthesis

Methyl 4-bromo-7-methyl-1*H*-indole-6-carboxylate **12**

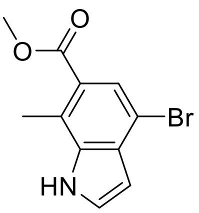

To a solution of methyl 5-bromo-2-methyl-3-nitrobenzoate **11** (500 mg, 1.82 mmol) in dry THF (18 ml) at −78 ˚C was added dropwise over 10 min a solution of vinylmagnesium bromide (1.0 M in THF, 5.84 mL, 5.84 mmol, 3.2 eq). The reaction mixture was stirred at −78°C for 1.5 h then allowed to warm to rt and stirred overnight. The reaction mixture was quenched by slow addition of $NH_4Cl$ (5 mL), concentrated *in vacuo*, resuspended in EtOAc (50 mL), then washed with $NH_4Cl$ (50 mL) and brine (2 × 50 mL), passed through a hydrophobic frit and concentrated *in vacuo*. Purification by FC ($SiO_2$, 5-40% EtOAc in pet ether 40-60) gave **12** (208 mg, 43%) as a cream coloured solid.

$^1H$ NMR (400 MHz, Chloroform-*d*) δ 8.51 (s, 1H), 7.96 (s, 1H), 7.43 (dd, *J* = 2.9, 2.9 Hz, 1H), 6.66 (dd, *J* = 3.2, 2.2 Hz, 1H), 3.94 (s, 3H), 2.78 (s, 3H). LCMS retention time = 2.20 min (100%), (m/z) $[M-H]^-$ = 266.

Methyl 7-methyl-4-(4,4,5,5-tetramethyl-1,3,2-dioxaborolan-2-yl)-1H-indole-6-carboxylate **13**

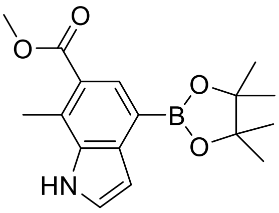

4-Bromo-7-methyl-indole-6-carboxylic acid methyl ester **12** (500 mg, 1.8 mmol, 1 equiv.), bis(pinacolato)diboron (585 mg, 2.3 mmol, 1.3 equiv.), potassium acetate (521 mg, 5.3 mmol, 3.0 equiv.) and Pd(dppf)$Cl_2$·DCM (145 mg, 0.18 mmol, 0.1 equiv.) were stirred in anhydrous DMSO (1 ml) and heated at 90˚C for 4 h, after which time the reaction was completed by LC-MS monitoring. The reaction mixture was allowed to cool to room temperature and water (10 mL) was added. The resulting precipitate was filtered and washed with water (10 mL). The organic residue was taken up in DCM (10 mL), washed with water (10 mL), brine (10 mL), dried (hydrophobic frit) and the solvent removed *in vacuo*. The crude material was purified by FCC (20% - 50% EtOAc/Pet ether) to yield an off-white solid 535 mg (96%).

$^1H$ NMR (400 MHz, $CDCl_3$) δ 8.37 (br s, 1H), 8.26 (s, 1H), 7.42 (t, *J* = 2.8 Hz, 1H), 7.10 (t, *J* = 2.8 Hz, 1H), 3.93 (s, 3H), 2.83 (s, 3H), 1.42 (s, 12H). LCMS retention time = 2.20 min (100%), (m/z) $[M-H]^-$ = 314.2, $[M+H]^+$ = 316.3.

2-(5-bromo-2-chlorophenoxy)pyridine **14**

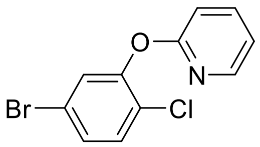

5-bromo-2-chlorophenol (312 mg, 1.50mmol) and 2-bromopyridine (359 mg, 2.28mmol) were reacted and purified according to Method D1, in dry DMF (1.3 ml) with $K_2CO_3$ at 150°C for 18h, to give the product **14** as a white solid (247 mg, 58%).

1H NMR (400 MHz, Acetone-d6) δ 8.10 (dd, J = 5.0, 1.6 Hz, 1H), 7.91 (ddd, J = 8.2, 7.2, 2.0 Hz, 1H), 7.55 – 7.46 (m, 3H), 7.18 – 7.10 (m, 2H). LCMS m/z 285.8 $(M+H)^+$.

4-(4-chloro-3-(pyridin-2-yloxy)phenyl)-7-methyl-1H-indole-6-carboxylic acid **16**

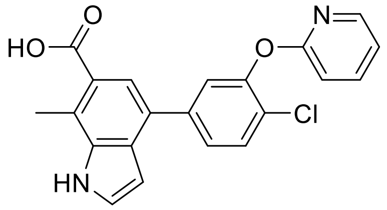

Methyl 4-(4,4,5,5-tetramethyl-1,3,2-dioxaborolan-2-yl)-1H-indole-6-carboxylate **13** (108 mg, 0.342 mmol) and 2-(5-bromo-2-chlorophenoxy)pyridine **14** (107 mg, 0.377 mmol) in DME:Water (3:1, 4 mL) were reacted and purified according to Method A to give methyl 4-(4-chloro-3-(pyridin-2-yloxy)phenyl)-7-methyl-1H-indole-6-carboxylate **15** (65 mg, 48%). LCMS m/z 393.2 $(M+H)^+$.

The methyl ester **15** (20 mg) was stirred with LiOH (11mg, 0.254 mmol, 5 equiv.) in THF:water (2:1, 1.1 mL) and reacted and purified according to Method B to give the product **16** as a white-off solid (3 mg, 16%).

$^1$H NMR (400 MHz, Acetone-d6) δ 10.81 (s, 1H), 8.14 (dt, J = 5.0, 1.3 Hz, 1H), 7.90 (t, J = 1.5 Hz, 1H), 7.88 (s, 1H), 7.69 (d, J = 8.8 Hz, 1H), 7.63 (tt, J = 5.4, 2.5 Hz, 3H), 7.19 – 7.10 (m, 2H), 6.76 (dd, J = 3.2, 1.8 Hz, 1H), 2.88 (s, 3H). LCMS retention time = 2.13 min (92%), (m/z) $[M-H]^-$ = 377.1, $[M+H]^+$ = 379.2.

4-(4-chloro-3-(pyridin-2-yloxy)phenyl)-N-(N,N-dimethylsulfamoyl)-7-methyl-1H-indole-6-carboxamide **CAM2602**

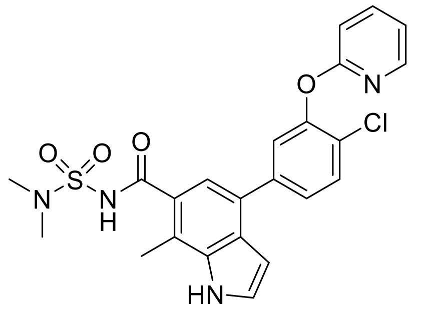

Acid **16** (18 mg, 0.05 mmol, 1.0 equiv.) and dimethylsulfamide (9 mg, 0.07 mmol, 1.5 equiv.) were reacted and purified according to Method F to give the product (5.4 mg, 22%) as a white solid.

$^1$H NMR (400 MHz, $CD_3OD$) δ 8.61 (s, 1H), 7.71-7.67 (m, 3H), 7.50 (s, *J* = 3.2 Hz, 1H), 7.32 (s, 1H), 7.09 (s, 1H), 6.70 (d, *J* = 3.2 Hz, 1H), 3.03 (s, 6H), 2.71 (s, 3H), 2.55 (s, 3H). LCMS retention time = 2.10 min (95%), (m/z) $[M-H]^-$ = 497.4, $[M+H]^+$ = 499.9.

Methyl 4-(4-chlorophenyl)-7-methyl-1*H*-indole-6-carboxylate **17**

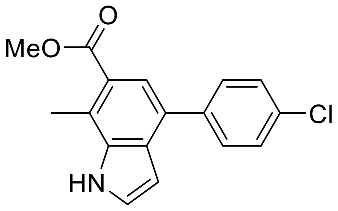

Compound **13** (220 mg, 0.82 mmol) and 4-chlorophenylboronic acid (154 mg, 0.98 mmol, 1.2 eq) were reacted according to Method A. Purification by FC ($SiO_2$, 8-66% EtOAc in pet ether 40-60) gave the product as a cream coloured solid (202 mg, 82%).

$^1$H NMR (400 MHz, Chloroform-*d*) δ 8.49 (s, 1H), 7.81 (s, 1H), 7.69 – 7.61 (m, 2H), 7.50 – 7.42 (m, 3H), 6.74 (dd, *J* = 3.2, 2.1 Hz, 1H), 3.95 (s, 3H), 2.85 (s, 3H). LCMS retention time = 2.46 min (100%), (m/z) [M-H]- = 298.

4-(4-Chlorophenyl)-7-methyl-1*H*-indole-6-carboxylic acid **4**

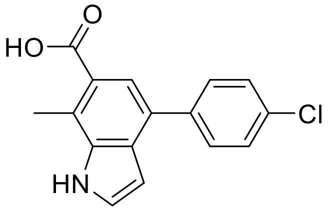

Methyl ester **17** (118 mg, 0.39 mmol) was deprotected with LiI (527 mg, 3.94 mmol, 10.0 eq) according to Method C. Purification by trituration with hexane gave the product as a buff solid (107 mg, 95%).
$^1$H NMR (400 MHz, Methanol-*d*$_4$) δ 7.73 (s, 1H), 7.71 – 7.63 (m, 2H), 7.54 – 7.45 (m, 3H), 6.65 (d, *J* = 3.2 Hz, 1H), 2.84 (s, 3H). LCMS retention time = 2.16 min (100%), (m/z) $[M-H]^-$ = 284.

4-(4-Chlorophenyl)-N-(cyclopropylsulfonyl)-7-methyl-1H-indole-6-carboxamide **7**

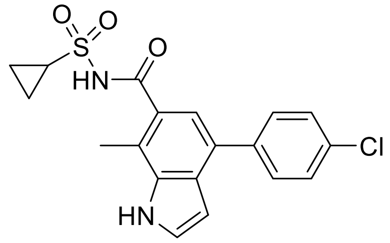

Compound **4** (40mg, 0.14 mmol, 1.0 eq) and cyclopropane sulfonamide (25 mg, 0.21 mmol, 1.5 eq) were reacted according to Method F. Purification by FC ($SiO_2$, 10-60% EtOAc in pet ether 40-60 (both with 0.5% AcOH)) gave a colourless oil which was dissolved in $Et_2O$, then precipitated with hexane to give the product as a white solid (40 mg, 73%).

$^1$H NMR (400 MHz, Methanol-*d*$_4$) δ 7.76 – 7.67 (m, 2H), 7.55 – 7.46 (m, 3H), 7.30 (s, 1H), 6.67 (d, *J* = 3.2 Hz, 1H), 3.21 (tt, *J* = 8.0, 4.8 Hz, 1H), 1.36 – 1.30 (m, 2H), 1.24 – 1.15 (m, 2H). LCMS retention time = 2.17 min (97%), (m/z) $[M-H]^-$ = 387.

4-(4-Chlorophenyl)-N-(N,N-dimethylsulfamoyl)-7-methyl-1H-indole-6-carboxamide **8**

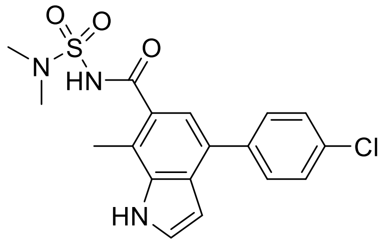

Compound **4** (80 mg, 0.27 mmol) and dimethylsulfamide (47 mg, 0.38 mmol) were reacted and purified according to Method F to give the product as a white solid. (50 mg, 47%).

$^1$H NMR (400 MHz, $CD_3OD$) δ 7.71 (d, *J* = 8.0 Hz, 2H), 7.50 (d, *J* = 8.0 Hz, 2H), 7.49 (s, 1H), 7.27 (s, 1H), 6.66 (d, *J* = 2.8 Hz, 1H), 3.04 (s, 6H), 2.70 (s, 3H). LCMS m/z 390.1 $(M-H)^-$

2-(5-bromo-2-chlorophenoxy)pyrazine **19**

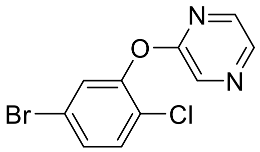

5-bromo-2-chlorophenol (500 mg, 2.41 mmol) and 2-fluoropyrazine (236 mg, 2.41 mmol) were reacted in dry DMSO (4 ml) with $CsCO_3$ at 90°C for 16 h according to Method D2. The crude was purified by FC ($SiO_2$, 0-20% EtOAc in Pet ether 40-60) to give the product as a white solid (444 mg, 65%).

$^1$H NMR (400 MHz, Acetone-d6) δ 8.60 (d, J = 1.4 Hz, 1H), 8.40 (d, J = 2.7 Hz, 1H), 8.14 (dd, J = 2.7, 1.4 Hz, 1H), 7.66 (dd, J = 1.9, 0.6 Hz, 1H), 7.56 (dd, J = 2.3, 1.3 Hz, 2H). LCMS retention time = 2.70 min (86%), (m/z) $[M+H]^+$ = 286.6.

4-(4-chloro-3-(pyrazin-2-yloxy)phenyl)-7-methyl-1H-indole-6-carboxylic acid **18**

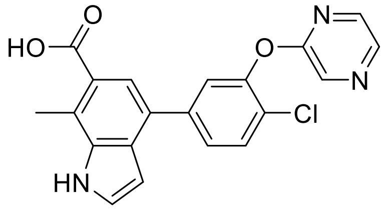

Methyl 4-(4,4,5,5-tetramethyl-1,3,2-dioxaborolan-2-yl)-1H-indole-6-carboxylate **13** (40 mg, 0.13 mmol) and 2-(5-bromo-2-chlorophenoxy)pyrazine **19** (42 mg, 0.15 mmol) were reacted in DME/Water=3:1 (2 ml) according to Method A to give methyl 4-(4-chloro-3-(pyrazin-2-yloxy)phenyl)-7-methyl-1H-indole-6-carboxylate as a white solid (44 mg, 84%). LCMS m/z 392.2 $(M-H)^-$. Methyl 4-(4-chloro-3-(pyrazin-2-yloxy)phenyl)-7-methyl-1H-indole-6-carboxylate (44 mg, 0.11 mmol) was reacted with LiOH (23 mg, 0.56 mmol, 5 equiv.) in THF/water=2:1 (2.25 ml) according to Method B to give the product as a white-off solid (7 mg, 16%).

$^1$H NMR (400 MHz, DMSO-d6) δ 12.54 (s, 1H), 11.64 (s, 1H), 8.70 (d, J = 1.4 Hz, 1H), 8.42 (d, J = 2.6 Hz, 1H), 8.23 (dd, J = 2.7, 1.4 Hz, 1H), 7.73 (d, J = 8.3 Hz, 1H), 7.67 – 7.59 (m, 4H), 6.63 (dd, J = 3.2, 1.8 Hz, 1H), 2.79 (s, 3H). LCMS retention time = 2.01 min (93%), (m/z) $[M-H]^-$ = 378.2, $[M+H]^+$ = 380.2.

4-(4-chloro-3-(pyrazin-2-yloxy)phenyl)-N-(N,N-dimethylsulfamoyl)-7-methyl-1H-indole-6-carboxamide **9**

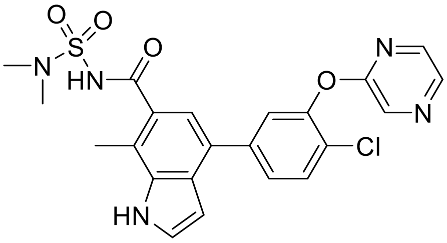

4-(4-chloro-3-(pyrazin-2-yloxy)phenyl)-7-methyl-1H-indole-6-carboxylic acid **18** (25 mg, 66 µmol), carbonyldiimidazole (32 mg, 0.198 mmol) and dimethylsulfamide (9 mg, 72 µmol) were reacted in THF (1.5 ml) according to Method F. The crude was purified by preparative HPLC (Column: Supelco Supelcosil® LC-18, 5-95% ACN in water + 0.1% formic acid) to give the product as a white solid (6 mg, 19%).

$^1$H NMR (400 MHz, Acetone-d6) δ 10.81 (s, 1H), 8.62 (d, J = 1.4 Hz, 1H), 8.39 (d, J = 2.6 Hz, 1H), 8.16 (dd, J = 2.7, 1.4 Hz, 1H), 7.78 – 7.75 (m, 1H), 7.74 – 7.69 (m, 2H), 7.64 – 7.59 (m, 1H), 7.51 (s, 1H), 6.80 – 6.72 (m, 1H), 3.02 (s, 6H), 2.74 (s, 3H). LCMS retention time = 2.10 min (100%), (m/z) $[M-H]^-$ = 484.2, $[M+H]^+$ = 486.2.

Methyl 4-(4-chloro-3-cyanophenyl)-7-methyl-1*H*-indole-6-carboxylate **20**

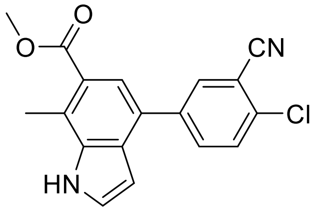

Compound **3** (240 mg, 0.90 mmol) and 4-chloro-3-cyanophenylboronic acid (211 mg, 1.16 mmol, 1.3 eq) were reacted and purified according to Method A. Purification by FC ($SiO_2$, 8-66% EtOAc in pet ether 40-60) gave the product as a cream coloured solid (186 mg, 64%).

$^1$H NMR (400 MHz, Chloroform-*d*) δ 8.58 (s, 1H), 8.01 (d, *J* = 2.2 Hz, 1H), 7.88 (dd, *J* = 8.4, 2.2 Hz, 1H), 7.80 (s, 1H), 7.64 (d, *J* = 8.4 Hz, 1H), 7.49 (dd, *J* = 2.9 Hz, 1H), 6.68 (dd, *J* = 3.2, 2.0 Hz, 1H), 3.97 (s, 3H), 2.87 (s, 3H). LCMS retention time = 2.33 min (100%), (m/z) [M-H]- = 323.

4-(4-Chloro-3-cyanophenyl)-7-methyl-1*H*-indole-6-carboxylic acid **6**

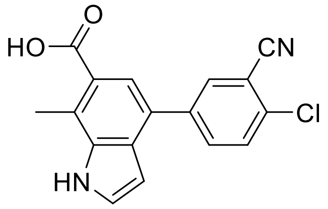

Methyl ester **20** (99 mg, 0.30 mmol) was deprotected according to Method C to give the product as a cream coloured solid (88 mg, 93%).

$^1$H NMR (400 MHz, Methanol-$d_4$) δ 8.09 (d, *J* = 2.2 Hz, 1H), 7.99 (dd, *J* = 8.5, 2.2 Hz, 1H), 7.80 – 7.73 (m, 2H), 7.55 (d, *J* = 3.1 Hz, 1H), 6.66 (d, *J* = 3.2 Hz, 1H), 2.86 (s, 3H). LCMS retention time = 2.04 min (98%), (m/z) [M-H]- = 309.

Methyl 4-(4-chloro-3-cyanophenyl)-1*H*-indole-6-carboxylate **21**

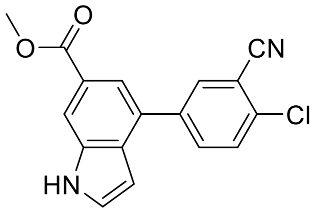

Methyl 4-bromo-1*H*-indol-6-carboxylate (100 mg, 0.39 mmol) and 4-chloro-3-cyanophenylboronic acid (79 mg, 0.43 mmol, 1.1 eq) were reacted according to Method A. Purification by FC ($SiO_2$, 8-66% EtOAc in pet ether 40-60), then trituration with $CH_2Cl_2$, gave the product as an off white solid (77 mg, 63%).

$^1$H NMR (400 MHz, Chloroform-*d*) δ 8.65 (s, 1H), 8.24 (d, *J* = 1.3 Hz, 1H), 8.04 (d, *J* = 2.1 Hz, 1H), 7.93 – 7.84 (m, 2H), 7.66 (d, *J* = 8.4 Hz, 1H), 7.51 (t, *J* = 2.9 Hz, 1H), 6.69 (t, *J* = 2.5 Hz, 1H), 3.99 (s, 3H). LCMS: retention time = 2.31 min (97%), m/z (ES−) 309 ([M−H]$^-$, 100%).

4-(4-Chloro-3-cyanophenyl)-1*H*-indole-6-carboxylic acid **9**

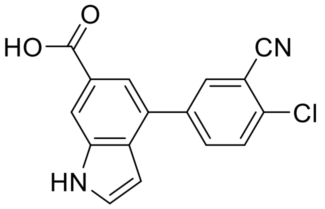

Methyl ester **21** (16 mg, 0.05 mmol) was hydrolysed with NaOH (6 mg, 0.15 mmol, 3 eq) and purified according to Method B to give the product as a cream coloured solid (12 mg, 79%).

NMR: $^1$H NMR (400 MHz, DMSO-$d_6$) δ 12.68 (s, 1H), 11.77 (s, 1H), 8.24 (d, *J* = 2.1 Hz, 1H), 8.13 (s, 1H), 8.05 (dd, *J* = 8.5, 2.2 Hz, 1H), 7.87 (d, *J* = 8.5 Hz, 1H), 7.77 – 7.69 (m, 2H), 6.67 (t, *J* = 2.3 Hz, 1H). LCMS: retention time = 2.02 min (98%), *m/z* (ES−) 295 ([M−H]$^-$, 100%).

Methyl 5-hydroxy-4'-(trifluoromethoxy)-[1,1'-biphenyl]-3-carboxylate **22**

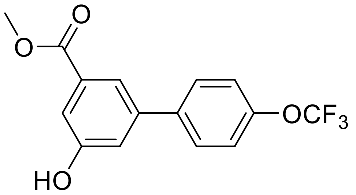

Methyl 3-hydroxy-5-(4,4,5,5-tetramethyl-1,3,2-dioxaborolan-2-yl)benzoate (42 mg, 0.15 mmol) and 1-bromo-4-(trifluoromethoxy)benzene (36 mg, 0.15 mmol) were reacted and purified according to Method A, to give the product as a white solid (37 mg, 78%).

$^1$H NMR (400 MHz, Chloroform-*d*) δ 7.84 (t, *J* = 1.5 Hz, 1H), 7.64 – 7.58 (m, 3H), 7.33 – 7.26 (m, 3H), 3.97 (s, 3H).

5-hydroxy-4'-(trifluoromethoxy)-[1,1'-biphenyl]-3-carboxylic acid **2**

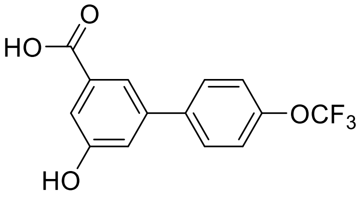

Methyl ester **22** (37 mg, 0.12 mmol) was hydrolysed with LiOH monohydrate (15 mg, 0.36 mmol) and purified according to Method B to give the product as a white solid (24 mg, 68%).

$^1$H NMR (400 MHz, Methanol-$d_4$) δ 7.76 (t, *J* = 1.6 Hz, 1H), 7.74 – 7.70 (m, 2H), 7.47 (dd, *J* = 2.4, 1.4 Hz, 1H), 7.38 (d, *J* = 8.2 Hz, 2H), 7.27 (t, *J* = 2.1 Hz, 1H). LCMS m/z 297.1 $(M\text{-}H)^-$

Methyl 4-(4-chlorophenyl)-1*H*-indole-6-carboxylate **23**

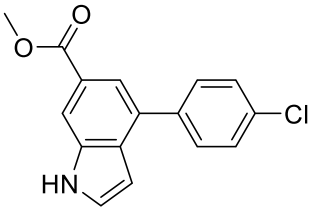

Methyl 4-bromo-1*H*-indol-6-carboxylate (200 mg, 0.79 mmol) and 4-chlorophenylboronic acid (148 mg, 0.94 mmol, 1.2 eq) were reacted according to Method A. Purification by FC ($SiO_2$, 6-50% EtOAc in pet ether 40-60) gave the product as a pale yellow solid (193 mg, 86%).

$^1$H NMR (400 MHz, Chloroform-*d*) δ 8.56 (s, 1H), 8.19 (d, *J* = 1.2 Hz, 1H), 7.88 (d, *J* = 1.3 Hz, 1H), 7.71 – 7.62 (m, 2H), 7.53 – 7.43 (m, 3H), 6.74 (t, *J* = 2.9 Hz, 1H), 3.98 (s, 3H). LCMS retention time = 2.37 min (95%), *m/z* (ES−) 284 ($[M−H]^-$, 100%).

4-(4-Chlorophenyl)-1*H*-indole-6-carboxylic acid **3**

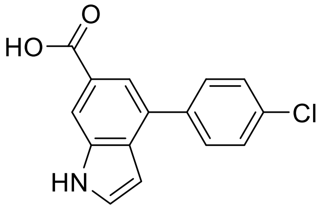

Methyl ester **23** (189 mg, 0.66 mmol) was hydrolysed with NaOH (79 mg, 1.98 mmol, 3 eq) according to Method B to give the product as a pale yellow solid (175 mg, 97%).

$^1$H NMR (400 MHz, Methanol-$d_4$) δ 8.17 (t, *J* = 1.2 Hz, 1H), 7.79 (d, *J* = 1.4 Hz, 1H), 7.75 – 7.66 (m, 2H), 7.56 – 7.48 (m, 3H), 6.67 (dd, *J* = 3.2, 0.9 Hz, 1H). LCMS retention time = 2.11 min (100%), *m/z* (ES−) 270 ($[M−H]^-$, 100%).
